# Structural mechanism of the type IX Retron-Kva2 anti-phage defense system

**DOI:** 10.64898/2026.07.14.738597

**Authors:** Yuya Hayashi, Yoshihisa Mitsuda, Kotaro Chihara, Kanta Yoneyama, Junichiro Ishikawa, Masahiro Hiraizumi, Masanori Hashino, Kazuhiro Horiba, Keitaro Yamashita, Kotaro Kiga, Hiroshi Nishimasu

**Author notes:** Corresponding authors, (K.K.) (H.N.). These authors contributed equally (Yuya Hayashi, Yoshihisa Mitsuda, Kotaro Chihara, Kanta Yoneyama, Junichiro Ishikawa).

## Abstract

Retrons are prokaryotic genetic elements that protect bacterial cells from invading phages. The type IX retron comprises a non-coding RNA, a reverse transcriptase (RT), and dual effectors: a putative HEPN nuclease and a winged helix-turn-helix (WH) protein. Here, we show that the type IX retron system from *Klebsiella variicola* cleaves host rRNAs and tRNAs upon phage infection, mediating anti-phage defense via an abortive-infection mechanism. Cryo-electron microscopy analysis reveals that the RT, HEPN, and WH proteins, along with multicopy single-stranded DNA (msDNA), form a unique, sheet-like supermolecular complex. Notably, the HEPN active site is encircled by the WH and msDNA within the complex, suppressing the nuclease activity prior to phage infection. A phage-encoded exonuclease cleaves the msDNA, likely releasing the HEPN nuclease to cleave host RNAs and induce growth arrest in infected cells. Overall, these findings highlight the structural and functional diversity of prokaryotic anti-phage defense systems.

## Introduction

Prokaryotes are constantly challenged by invading mobile genetic elements, particularly bacteriophages, and have evolved a broad repertoire of defense systems to survive these infections^1–4^. Some defense systems, such as restriction-modification systems and CRISPR-Cas adaptive immunity, directly recognize and cleave foreign nucleic acids via sequence-specific or RNA-guided nuclease activities^5,6^. Others employ indirect strategies, such as abortive infection, in which infected cells undergo growth arrest or cell death before phages complete their replication cycle, thereby limiting phage propagation within the bacterial population^7^.

Retrons are prokaryotic genetic elements originally identified by their production of multicopy single-stranded DNA (msDNA), an unusual RNA–DNA hybrid synthesized by a retron-encoded reverse transcriptase (RT)^8–10^. A typical retron locus encodes a non-coding RNA (ncRNA), an RT, and one or more associated effector proteins. The ncRNA contains *msr* and *msd* regions (referred to as msrRNA and msdRNA, respectively). During msDNA synthesis, the RT reverse-transcribes the *msd* region of the ncRNA (msdRNA) to produce a complementary DNA strand (msdDNA) that is covalently linked to the msrRNA via a 2′–5′ bond formed at the 2′-OH group of a conserved guanosine (the branching guanosine)^9–11^. The RNA template is then degraded by the host RNase H, yielding the characteristic RNA–DNA hybrid.

Although retrons were discovered several decades ago, their physiological roles remained unclear for a long time. However, recent studies revealed that many retrons function as anti-phage defense systems^12–14^. In these systems, the RT–msDNA complex associates with cognate effector proteins to form an antitoxin–toxin complex: the RT–msDNA complex acts as an antitoxin to suppress the effector’s toxicity under uninfected conditions, whereas phage infection relieves this inhibition and activates the effector, thereby facilitating abortive infection. Retron systems are diverse and classified into 13 distinct types, based on their effector proteins^15,16^. Subsequent studies have begun to reveal the remarkable functional and mechanistic diversity among retron systems. For example, in the type II-A1 Retron-Eco1 system (formerly Retron-Ec86)^17^, the RT–msDNA complex associates with an *N*-glycosidase effector to form a filament-like assembly that facilitates NAD(P)^+^ depletion^18–20^. Retron-Eco4 (formerly Retron-Ec83)^21^ and Retron-Eco7 (formerly Retron-Ec78)^22^ from the type I-A systems encode dual PtuA–PtuB helicase-nuclease effectors that cleave nucleic-acid substrates, including single-stranded DNA^23^ and specific tRNAs^24–29^. The type I-B2 Retron-Eco8 system employs an OLD-family nuclease effector to degrade genomic DNA^30–33^, while the type I-C1 Retron-Eco2 system uses a TOPRIM nuclease to cleave tRNA substrates^34,35^.

The type IX retron system encodes an RT, an ncRNA, and two predicted effector proteins: a putative HEPN-family nuclease and a winged helix–turn–helix-like (WH) protein^15,16^. HEPN nucleases have RNase activities and are found in various prokaryotic defense and toxin systems^36^, but the function of the HEPN effector in the type IX retrons remains unknown. Intriguingly, the type IX HEPN effector lacks the canonical Rx_4–6_H motif conserved among typical HEPN nucleases, and instead contains conserved His and Arg residues (H64 and R122 in the HEPN effector of the *Klebsiella variicola* retron). The role of the WH protein is also unclear, as most retron systems encode a single effector enzyme along with an RT–ncRNA module.

In this study, we combine genetic, biochemical, and structural analyses to elucidate the anti-phage defense mechanism of the type IX retron from *K. variicola* (referred to as Retron-Kva2). We show that the RT–msDNA subcomplex associates with the HEPN–WH subcomplex to form a sheet-like supermolecular assembly that suppresses the HEPN’s RNA-cleavage activity. Our data further suggest that phage-derived nucleases trigger the disassembly of the Retron-Kva2 complex, enabling the HEPN nuclease to cleave rRNAs and tRNAs and thereby induce abortive infection via growth arrest. Overall, these findings provide mechanistic insight into the type IX retron system and broaden our understanding of the functional and mechanistic diversity of retron-mediated immunity.

## Results

### Anti-phage defense activity of Retron-Kva2

To investigate the function of the type IX retron, we expressed the *K. variicola* Retron-Kva2 system (HEPN, WH, ncRNA, and RT) in *Escherichia coli*, infected the bacterial cells with 81 different phages from both the BASEL collection and our in-house phage library, and then examined their anti-phage defense activities using spot assays (Figures 1A and 1B). Retron-Kva2 protected *E. coli* from phages of several families, including *Demerecviridae*, *Vequintavirinae*, and *Queuovirinae* (Figure 1B). The D223A/D224A mutations in the RT’s YADD catalytic motif abolished the defense activity (Figures 1C and S1A), confirming the functional importance of the RT-mediated msDNA production. Mutations in the putative catalytic residues in the HEPN gene (H64A and R122A) also abolished the defense activity (Figures 1C and S1A), indicating that both H64 and R122 function as catalytic residues and that HEPN’s nuclease activity is essential for anti-phage defense by Retron-Kva2. Furthermore, the deletion of the WH gene abolished the defense activity (Figures 1C and S1A), highlighting the requirement of the WH protein for Retron-Kva2 function. The expression of HEPN, but not its mutants (H64A and R122A), inhibited bacterial growth even in the absence of phage infection (Figure 1D), indicating that HEPN’s nuclease activity is essential for growth inhibition. Co-expression with RT–ncRNA–WH completely suppressed HEPN’s toxicity, while co-expression with WH alone also effectively mitigated this toxicity (Figure 1D). These results suggest that HEPN functions as the toxin and RT–msDNA–WH acts as the antitoxin, while WH alone can also act as an antitoxin. We next monitored the growth of *E. coli* cells containing Retron-Kva2 after infection with the Bas24 phage. *E. coli* cells with Retron-Kva2 exhibited normal growth under low multiplicity of infection (MOI) conditions, whereas their growth was inhibited under high MOI conditions (Figures 1E and S1B). Together, these results suggest that Retron-Kva2 is activated by phage infection and induces growth arrest in the infected cells, thereby protecting the bacterial population from phage infection through an abortive infection mechanism.

**Figure 1.**
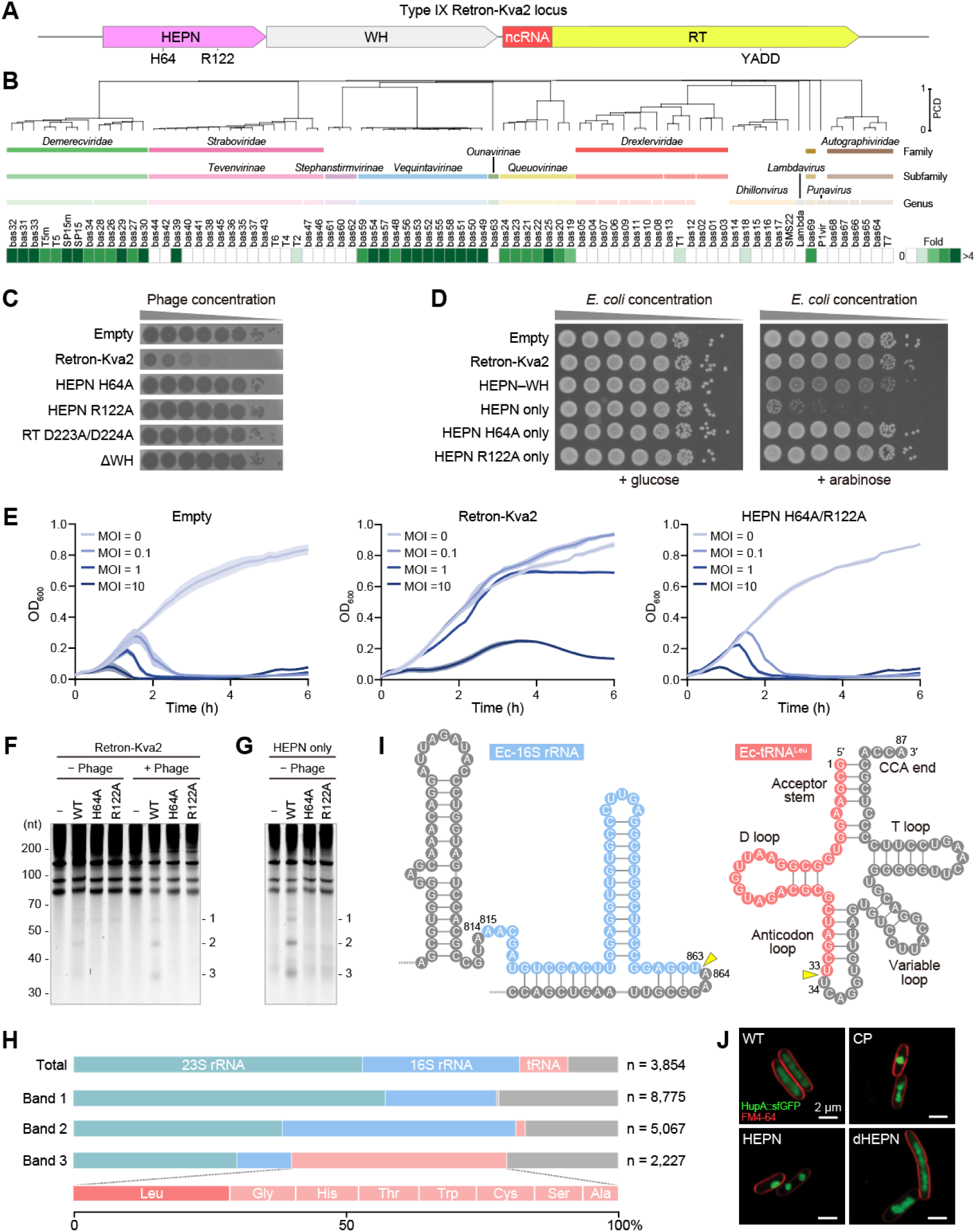
Function of Retron-Kva2. (**A**) Schematic of the Retron-Kva2 operon. (**B**) Defense profile of Retron-Kva2 against BASEL phage collections and common laboratory coliphages. The phage order and dendrogram were determined through hierarchical clustering based on the proteome composition distance matrix. (**C**) Anti-phage defense activities of wild-type (WT) Retron-Kva2, Retron-Kva2 with the HEPN mutations (H64A or R122A), Retron-Kva2 with the RT mutations (D223A/D224A), and Retron-Kva2 with the WH deletion against the Bas24 phage, evaluated by spot assays. *E. coli* cells transformed with an empty vector were used as the control. Assays were repeated at least three times with similar results. (**D**) Growth of *E. coli* cells harboring plasmids encoding either Retron-Kva2, HEPN–WH, HEPN, or HEPN mutants (H64A or R122A), in the presence of 0.2% glucose (repressive condition) or 0.2% arabinose (inductive condition). *E. coli* cells transformed with an empty vector were used as a control. Assays were repeated at least three times with similar results. (**E**) Growth curves of *E. coli* cells harboring an empty vector (−) or plasmids expressing Retron-Kva2 with or without the HEPN mutations (H64A/R122A). The cells were infected with the Bas24 phage at a multiplicity of infection (MOI) of 0, 0.1, 1, or 10. Data represent the mean ± SD of three biological replicates (n = 3). (**F**) RNA cleavage by Retron-Kva2. *E. coli* cells were transformed with an empty vector (−) or plasmids expressing either WT Retron-Kva2 or Retron-Kva2 with HEPN mutations (H64A or R122A), and were grown with or without T5 phage infection. RNA was extracted from the cells 1.5 h after phage infection. RNA was fractionated by urea-PAGE, and the gels were stained with SYBR Gold. Positions for RNA cleavage products (bands 1–3) are indicated. Assays were repeated at least three times with similar results. (**G**) RNA cleavage by HEPN. *E. coli* cells were transformed with an empty vector (−) or plasmids expressing HEPN alone (WT, H64A, and R122A), and RNA was extracted from the cells 1 h after HEPN induction. RNA was fractionated by urea-PAGE, and the gels were stained with SYBR Gold. Positions for RNA cleavage products (bands 1–3) are indicated. Assays were repeated at least three times with similar results. (**H**) Composition of the RNA cleavage products. The entire region encompassing the three major RNA cleavage products (bands 1–3) in (**F**) was purified from the gel, and their sequences were determined by Nanopore sequencing. Total, sum of bands 1–3. (**I**) Nucleotide sequences of *E. coli* 16S rRNA and tRNA^Leu^. Identified cleavage fragments are highlighted with different colors (blue for 16S rRNA and red for tRNA^Leu^), with the cleavage sites indicated by yellow triangles. RNA modifications are omitted for clarity. (**J**) Nucleoid morphology of *E. coli* cells expressing a chromosomally encoded *hupA*::sfGFP fusion. The cells were transformed with an empty vector or plasmids expressing HEPN or the H64A/R122A mutant (dHEPN), and were imaged by Airyscan confocal microscopy 60 min after HEPN induction. The cell boundaries were delineated with the membrane dye FM4-64. The cells harboring an empty vector, with or without chloramphenicol (CP) treatment (which induces nucleoid compaction), were used as controls.

### RNA cleavage by Retron-Kva2

Previous studies showed that the HEPN nucleases in CRISPR-Cas13 and HEPN-MNT toxin–antitoxin systems cleave various host RNAs, including tRNAs^37–39^ and rRNAs^40^, thereby facilitating anti-phage defense via abortive infection. Therefore, we investigated the RNA cleavage activity of the Retron-Kva2 system *in E. coli*. We infected *E. coli* cells—with and without plasmids expressing Retron-Kva2 or HEPN alone—with the T5 phage, extracted total RNA from the cells, and analyzed the RNA using denaturing urea-PAGE. Three major additional RNA bands (bands 1–3) were observed in the lysates from phage-infected bacterial cells expressing Retron-Kva2, but not from those lacking Retron-Kva2 or from those expressing Retron-Kva2 with the HEPN mutations (H64A or R122A) (Figure 1F). These additional bands were not detected in the lysates from uninfected cells (Figure 1F). Furthermore, the expression of HEPN alone, but not the HEPN mutants (H64A or R122A), resulted in the production of similar RNA bands (Figure 1G). These results indicate that Retron-Kva2 is activated upon phage infection and cleaves host RNAs using the HEPN effector.

To determine the identity of the cleavage products, we excised the region encompassing the three bands from the gel, purified the RNA fragments, and analyzed their nucleotide sequences using Nanopore sequencing. Our analysis revealed that the mapped reads were primarily derived from *E. coli* 23S rRNA (53.1%), 16S rRNA (28.9%), and tRNA (8.8%), while other fractions (9.2%) contained 5S rRNA and mRNA (Figure 1H). Bands 1 and 2 were predominantly composed of rRNA-derived fragments (23S rRNA (57.2%) and 16S rRNA (20.5%) for band 1, and 23S rRNA (38.3%) and 16S rRNA (42.9%) for band 2) (Figure 1H). In contrast, band 3 contained abundant fragments derived from various tRNAs (52.2%), including tRNA^Leu^ (10.7%) (Figure 1H). Among these, the fragments derived from 16S rRNA (nucleotides A815–U863) (6.6% in band 2) and tRNA^Leu^ (nucleotides G1–U33) (3.9% in band 3) were the most abundant (Figure 1I). Both RNAs share identical sequences between *K. variicola* and *E. coli*, indicating that Retron-Kva2 cleaves these RNA species in the natural host. Analysis of the 3′ ends of the tRNA-derived fragments revealed an enrichment of uracil, indicating that Retron-Kva2 HEPN preferentially cleaves the tRNA substrates between U and N (5′-U|N-3′) at the anticodon loop (Figures S2A and S2B), consistent with the preferences of several CRISPR-Cas13 nucleases^39^. Uracil was also enriched at the 3′ ends of the 16S rRNA-derived fragments (Figure S2C). In contrast, both ends of the 23S rRNA-derived fragments and the 5′ ends of the 16S rRNA-derived fragments exhibited a preference for A rather than U (Figures S2D–S2F), suggesting degradation by other nucleases after HEPN-mediated cleavage. Together, these results indicate that Retron-Kva2 cleaves rRNAs and tRNAs upon phage infection, thereby inducing translation arrest.

To examine the downstream consequences of RNA cleavage by Retron-Kva2, we expressed either WT HEPN or the HEPN H64A/R122A mutant in the *E. coli hupA*::sfGFP reporter strain and monitored the nucleoid morphology. The expression of WT HEPN, but not the HEPN mutant, induced nucleoid compaction, a hallmark of translation arrest^41^, as observed in the cells treated with chloramphenicol (Figure 1J), confirming that translation arrest is induced by the HEPN-mediated cleavage of rRNAs and tRNAs. Collectively, these findings indicate that the Retron-Kva2 system is activated upon phage infection, cleaving essential components of the translational machinery and inducing growth arrest in infected cells.

### Cryo-EM structure of the Retron-Kva2 complex

To elucidate the action mechanism of Retron-Kva2, we aimed to determine the cryo-EM structure of the Retron-Kva2 complex. We co-expressed three protein components—Strep-tagged RT, WH, and HEPN H64A—along with the ncRNA in *E. coli*, as we were unable to purify the Retron-Kva2 complex containing the WT HEPN effector due to poor expression, likely caused by its toxicity. We purified the Retron-Kva2 complex (with the HEPN H64A mutation) using Strep-Tactin affinity and size-exclusion chromatography (Figure S3A). SDS-PAGE and urea-PAGE analyses of the peak fraction from size-exclusion chromatography revealed that the RT, WH, and HEPN proteins form a complex with ∼40-nt and ∼60-nt nucleic acids (Figures S3B and S3C). Urea-PAGE analysis of the complex after DNase or RNase treatment confirmed that the bound nucleic acids represent msDNA, consisting of msrRNA (∼40 nt) and msdDNA (∼65 nt) (Figure S3C), consistent with a previous study^42^. Sequencing analysis indicated that nucleotides G45–A115 in the ncRNA (msdRNA) were primarily reverse-transcribed to produce the msdDNA (Figure S3D). These results suggest that (1) unlike typical retron systems, G139 and G121 at the end of stem loop 3 serve as the branching and opposing guanosine residues, respectively, and (2) nucleotides G45–C120 in the ncRNA are primarily reverse-transcribed into the msdDNA (nucleotides dG1–dC76), which is subsequently processed between dA5 and dT6 by exonuclease VII in *E. coli*, as reported in other retron systems^43^ (Figure S4A and S4B).

We determined the cryo-EM structure of the RT–msDNA–HEPN–WH complex (referred to as the Retron-Kva2 complex) at 2.9-Å resolution (Figures 2A–2D, S5A–H, S6, Table S1, and Video S1). The monomeric unit of the Retron-Kva2 complex consists of an RT–msDNA subcomplex and a HEPN–WH subcomplex, with the WH subunit interacting with both the RT–msDNA module and the HEPN effector. The C-terminal and middle domains of the WH subunit in one monomeric unit interact with the stem-loop region of the msDNA and the middle domain of the WH subunit in the adjacent monomeric unit, respectively, forming a head-to-tail dimer (Figures 2B–2D). Intriguingly, the HEPN subunits in one dimer extensively interact with the HEPN subunits in adjacent dimers, forming a higher-order, sheet-like assembly (Figures 2B–2D). Notably, the active site in the HEPN dimer is surrounded by the WH subunit and the msdDNA (Figures 2E and 2F), indicating that RNA substrates cannot access the HEPN active site within the Retron-Kva2 complex. Together, the cryo-EM structure reveals that RT–msDNA and WH associate with the HEPN effector to suppress its nuclease activity by sterically preventing RNA substrates from reaching the HEPN active site within the Retron-Kva2 complex.

**Figure 2.**
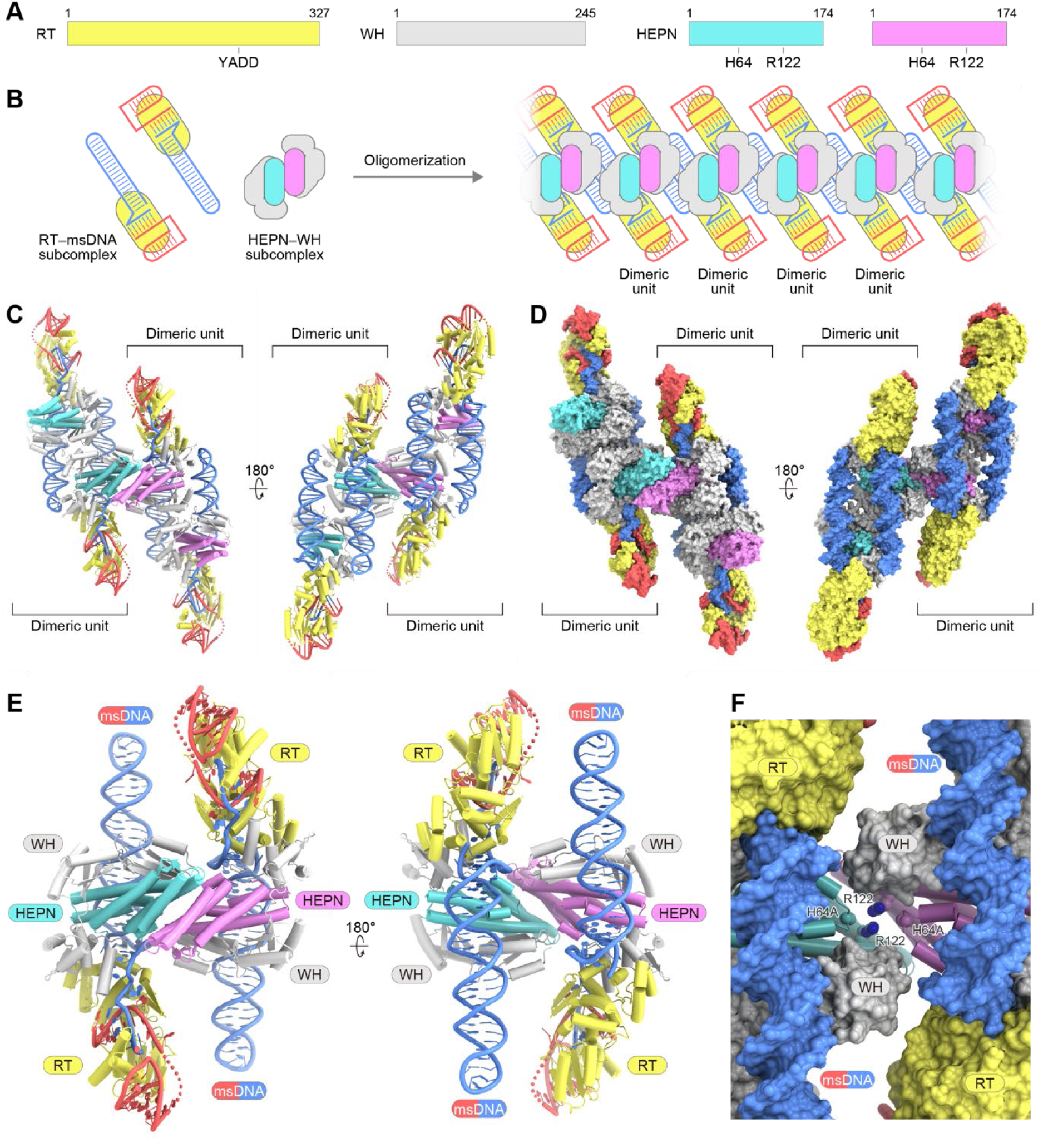
Structure of the Retron-Kva2 complex. (**A**) Schematic of the protein components of Retron-Kva2. (**B**) Oligomeric assembly of the Retron-Kva2 complex. (**C** and **D**) Cartoon (**C**) and surface (**D**) representations of two dimeric units of the Retron-Kva2 complex. (**E**) Cartoon representation of the Retron-Kva2 dimeric complex. (**F**) Close-up view of the active site in the HEPN dimer. The HEPN subunits and other components are shown as cartoon and surface representations, respectively.

### Structure of the RT–msDNA subcomplex

The structure reveals that the msDNA comprises a 35-nt msrRNA (G18–A52) and a 71-nt msdDNA (dT6–dC76), with G45–A51 in the msrRNA and dA70–dC76 in the msdDNA forming an RNA–DNA heteroduplex bound to the RT active site (Figures 3A, 3B, S7A, and S7B). Cryo-EM density was detected for dT6 but not for dG1–dA5 (Figure S7C), consistent with our sequencing data indicating that the msdDNA was processed by exonuclease VII (Figure S3D). No densities were observed for G1–A17, U37–U40, and G139–C173 in the msrRNA (Figures 3A and S7C), suggesting that these regions are disordered due to their flexibility. The msrRNA contains a stem loop (G18–C34) and a single-stranded linker region (nucleotides A35–G44), while the msdDNA contains a stem loop (dT6–dA63) and a single-stranded linker region (dT64–dT69) (Figures 3A, S7B, and S7C).

**Figure 3.**
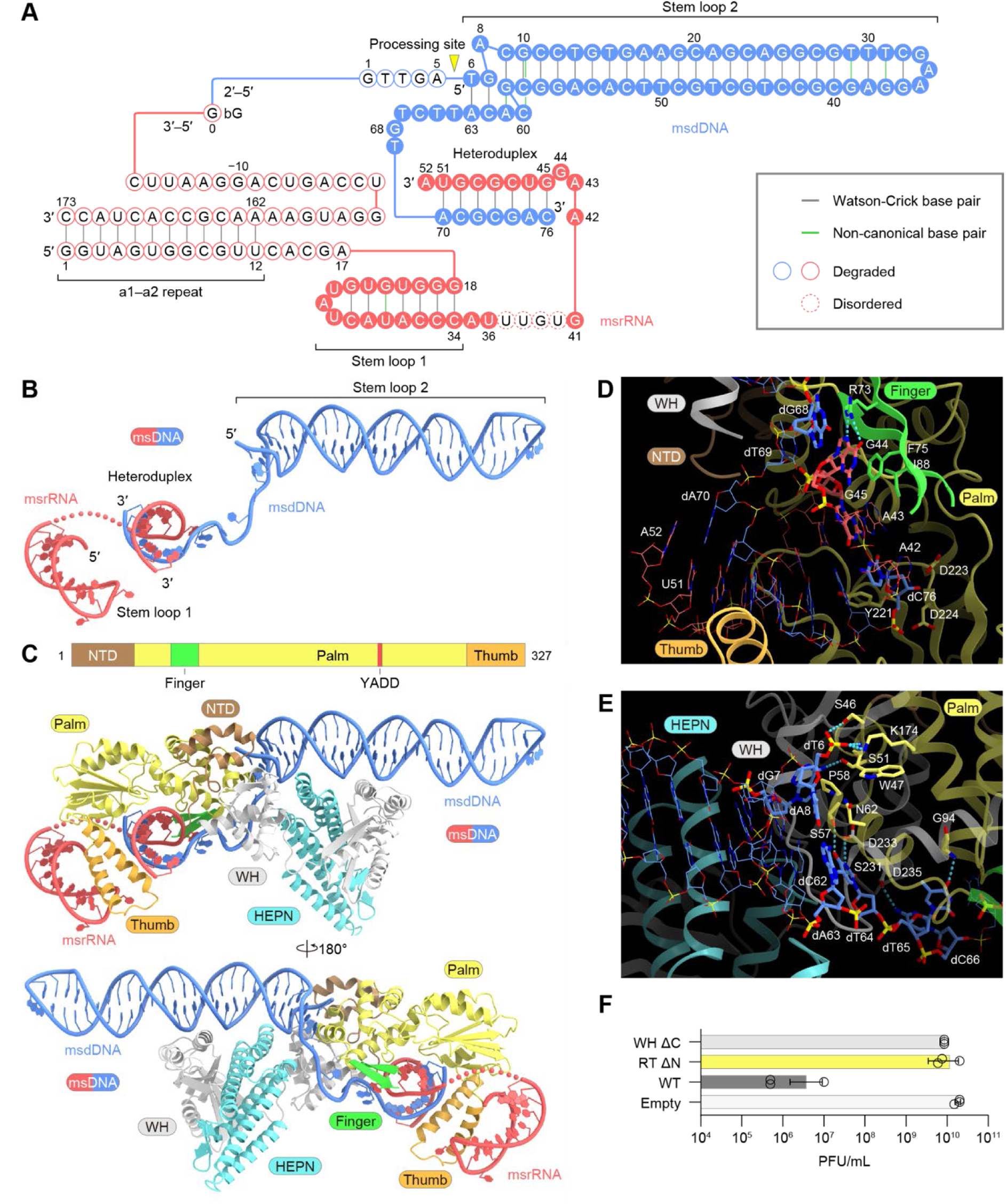
Structure of the Retron-Kva2 monomeric complex. (**A**) Nucleotide sequence of the msDNA. The ExoVII cleavage site is indicated by a yellow triangle. Watson–Crick and non-canonical base pairs are shown with gray and green lines, respectively. bG, branching G. (**B**) Structure of the msDNA. (**C**) Structure of the Retron-Kva2 monomeric complex (the RT–msDNA subcomplex bound to the HEPN–WH subcomplex). The domain arrangement of the RT is shown above the structure. In (**B**) and (**C**), disordered nucleotides between the heteroduplex and stem loop 1 are indicated by dotted lines. (**D**) Close-up view of the RT active site. Hydrogen bonds are depicted by cyan dashed lines. (**E**) Recognition of the msdDNA stem loop. (**F**) Anti-phage defense activities of WT Retron-Kva2 and its mutants against the Bas24 phage. The defense activities (PFU/mL) were evaluated by spot assays. *E. coli* cells transformed with an empty vector were used as the control. Data represent the mean ± SD of four independent replicates (n = 4). WH ΔC, Retron-Kva2 lacking the C-terminal 14 residues of the WH subunit; RT ΔN, Retron-Kva2 lacking the N-terminal 39 residues of the RT subunit.

Like the RT enzymes in other retron systems, such as Retron-Eco7^28^, the Retron-Kva2 RT adopts a typical right-hand-like fold comprising palm, finger, and thumb domains (Figures 3C and S8A). Similar to the Retron-Eco7 complex, the msrRNA stem loop is recognized by the thumb domain, while the heteroduplex is accommodated within the active site formed by the palm, finger, and thumb domains, with the conserved YADD motif (residues 221–224) located near the msdDNA 3′ end (Figures 3C, 3D, and S9). Notably, the incoming nucleotide-binding site is occupied by A42 in the msrRNA linker, while the template nucleotide G44 is flipped out from the heteroduplex and hydrogen bonds with R73, with its nucleobase stacked between F75 and dG68 (Figures 3D and S9). Thus, this Retron-Kva2 structure represents a reverse-transcription termination state, stabilized by interactions between the msrRNA stem loop and the RT’s thumb domain, as seen in other retron complexes^19,28^.

Unlike the Retron-Eco7 RT^28^, the Retron-Kva2 RT possesses an additional N-terminal domain (residues 1–45), which interacts with the C-terminal domain of the WH subunit (Figures 3C and S8A). The msdDNA linker region binds to a groove formed by the WH’s C-terminal helix and the RT’s palm/finger domains (Figures 3C, 3D, and S9). The basal region of the msdDNA stem loop adopts a distinctive structure, containing the flipped-out dA8 and two successive base triples (C9-G59-A61 and G10-C58-C60) (Figure S7B). This region is recognized by the RT and WH subunits, contributing to the orientation of the stem loop (Figures 3E and S9). The orientations of the msdDNA stem loops relative to the RT subunits differ substantially between the Retron-Kva2 and Retron-Eco7 complexes, primarily due to these distinct interactions with their msdDNAs (Figure S8A). The anti-phage defense activity of Retron-Kva2 was abolished by the deletion of either the RT’s N-terminal 39 residues or the WH’s C-terminal 14 residues (Figure 3F), confirming that the RT–msDNA–WH interactions are functionally essential.

### Structure of the HEPN–WH subcomplex

HEPN comprises five core α-helices and a small three-stranded β-sheet, resembling other HEPN nucleases, such as that in the HEPN-MNT toxin–antitoxin system^37^ (PDB 7AE8; RMSD = 4.4 Å for 127 equivalent Cα atoms), despite having limited sequence identity (9%) (Figures 4A and S8B). In the Retron-Kva2 complex, the HEPN subunits from one dimeric unit extensively interact with those from adjacent units, forming a dimer similar to typical HEPN nucleases (Figure 4A). As previously described^16^, the Retron-Kva2 HEPN protein lacks the canonical Rx_4–6_H catalytic motif but contains conserved H64 and R122 residues on the third and fourth core α-helices, respectively, creating an active site at the dimer interface (Figures 4A and S8B), consistent with our data showing the functional importance of both H64 and R122 (Figures 1C–1F, S1A, and S1B). Intriguingly, H64 and R122 from two different protomers of Retron-Kva2 HEPN occupy positions similar to the His (H107) and Arg (R102) residues in each protomer of HEPN–MNT (Figure S8B), highlighting the structural divergence among HEPN nucleases.

**Figure 4.**
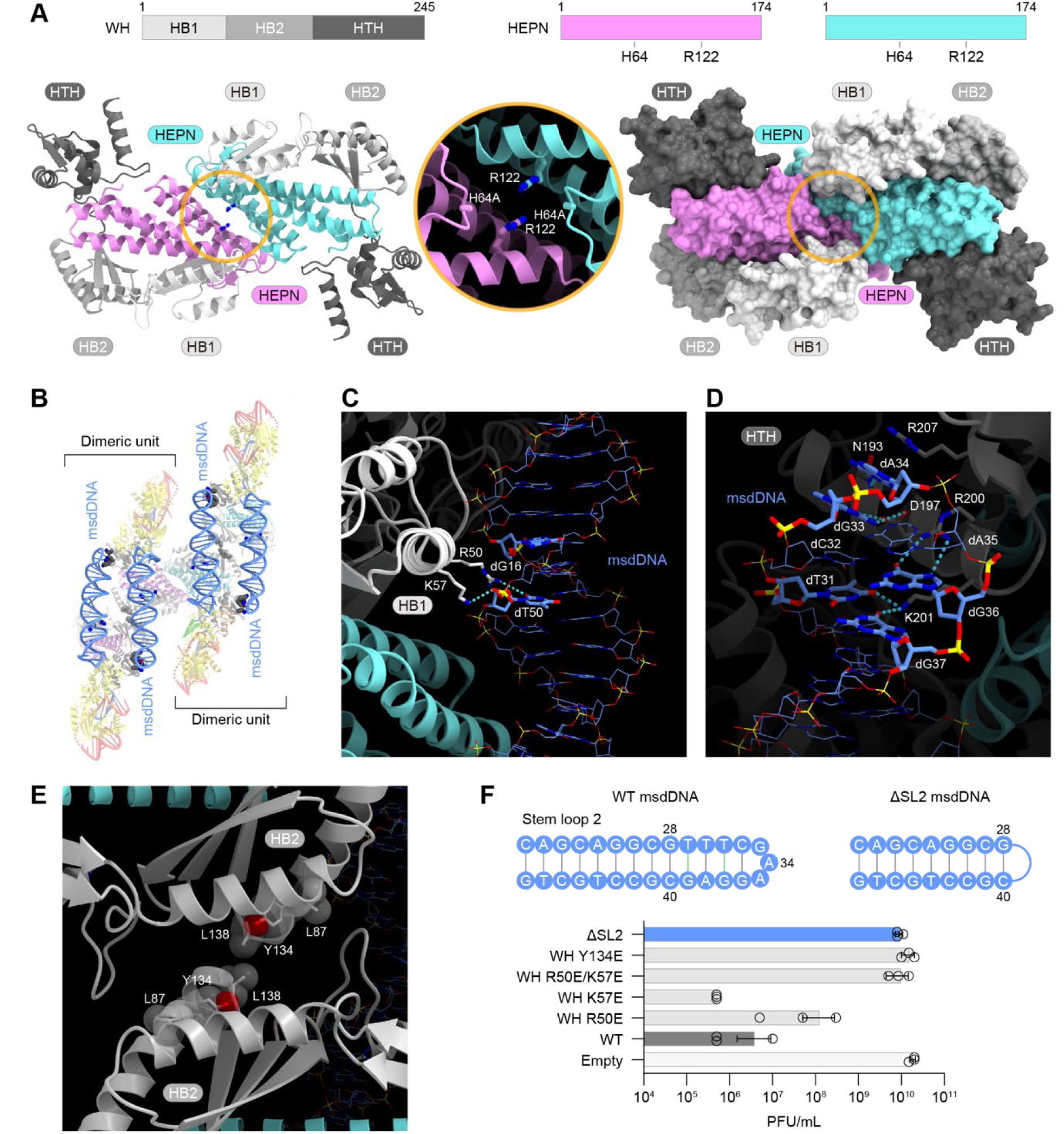
Oligomeric assembly of the Retron-Kva2 complex. (**A**) Structures of the HEPN–WH subcomplexes from two dimeric units. Domain arrangements are shown above the structures, in ribbon (left) and surface (right) representations. A close-up view of the HEPN active site is shown at the center. (**B**) WH-mediated interactions. Two dimeric units are shown in a semi-transparent ribbon representation to highlight msdDNA stem-loop 2. WH residues that interact with the msdDNA are shown as space-filling models. (**C** and **D**) Recognition of the middle (**C**) and loop (**D**) regions of msdDNA stem-loop 2. Hydrogen bonds are depicted by cyan dashed lines. (**E**) Interactions between adjacent WH subunits within a dimeric unit. (**F**) Anti-phage defense activities of WT Retron-Kva2 and its mutants against the Bas24 phage, evaluated by spot assays. Data represent the mean ± SD of four independent replicates (n = 4). ΔSL2, Retron-Kva2 lacking nucleotides dT29–dG39 of the msdDNA stem loop 2.

WH consists of three domains: domain 1 (residues 1–72), domain 2 (residues 73–147), and domain 3 (residues 148–245) (Figures 4A and S8C). Domains 1 and 2 share 17% sequence identity and adopt similar αββαβ folds (RMSD = 2.3 Å for 55 equivalent Cα atoms) to those found in various proteins such as the cytoplasmic domain of the zinc transporter^44^ (PDB 5VRF; RMSD = 3.1 Å for 64 equivalent Cα atoms, 5% sequence identity) (Figure S8C). Domain 3 contains five α-helices and a small β-hairpin, and resembles the HTH domain of the *N*-glycosidase effector in the Retron-Eco1 system^19,20^ (PDB 7XJG; RMSD = 2.2 Å for 91 equivalent Cα atoms, 20% sequence identity) (Figure S8C). The three domains of the WH subunit wrap around the HEPN subunit, with domain 1 located near the HEPN active site (Figure 4A). We refer to domains 1 and 2 as HEPN-binding domains 1 and 2 (HB1 and HB2), respectively, and domain 3 as the HTH domain.

### Oligomeric assembly of the Retron-Kva2 complex

The HEPN–WH subcomplex associates with the RT–msDNA subcomplex primarily through the interactions between the WH HTH domain and the RT N-terminal region (as stated above) and between the WH HB1 domain and the middle region of the double-stranded msdDNA (Figure 4B). R50 inserts into the minor groove side of the msdDNA and contacts dG16, while K57 interacts with the dT50 backbone phosphate (Figures 4C and S9). Furthermore, two RT–msDNA–HEPN–WH complexes form a dimeric assembly through interactions between the msdDNA stem loop and the WH HTH domain and between the WH HB2 domains (Figure 4B). Notably, dT31, dG33, dA34, and dG36 in the msdDNA stem loop form hydrogen bonds with K201, D197, N193, and R200 in the WH HTH domain, respectively, while dA34 stacks with R207 (Figures 4D and S9). Meanwhile, L87, Y134, and L138 in the WH HB2 domain of one complex hydrophobically interact with the equivalent residues in its adjacent complex (Figure 4E). The mutations of these WH residues and the deletion of nucleotides dT29–dG39 in the msdDNA stem loop reduced or abolished the anti-phage defense by Retron-Kva2 (Figure 4F). Together, these results highlight the importance of the dimeric assembly in the Retron-Kva2 complex for suppressing the toxicity of the HEPN effector and thereby facilitating the anti-phage defense.

### Retron-Kva2 activation by a phage-encoded exonuclease

To clarify how Retron-Kva2 detects phage infection, we infected *E. coli* carrying Retron-Kva2 with the Bas24 phage and isolated several escaper phages that evaded retron-mediated immunity (Figure 5A). Whole-genome sequencing of the escapers revealed point mutations in various genes, including one encoding a putative exonuclease (referred to as Bas24-Exo) (Figure 5B and Table S2). An AlphaFold3 prediction indicated that Bas24-Exo is structurally similar to known exonucleases, such as λ exonuclease^45^ (PDB 3SLP; RMSD = 2.8 Å for 169 equivalent Cα atoms, 19% sequence identity) (Figure 5C). Indeed, the purified Bas24-Exo protein efficiently cleaved single-stranded DNA but not double-stranded DNA *in vitro* (Figure 5D), confirming its nuclease activity. The escaper phages carry an E197V point mutation in the Bas24-Exo gene, and the purified Bas24-Exo E197V mutant exhibited reduced DNA-cleavage activity as compared to WT (Figure 5D). Consistent with this reduced activity, the E197V mutation is located near the active site in the predicted structure of Bas24-Exo (Figure 5C). Furthermore, the co-expression of WT Bas24-Exo, but not the E197V mutant, caused growth defects in *E. coli* containing Retron-Kva2 (Figure 5E), implicating the nuclease activity of Bas24-Exo in Retron-Kva2 activation.

**Figure 5.**
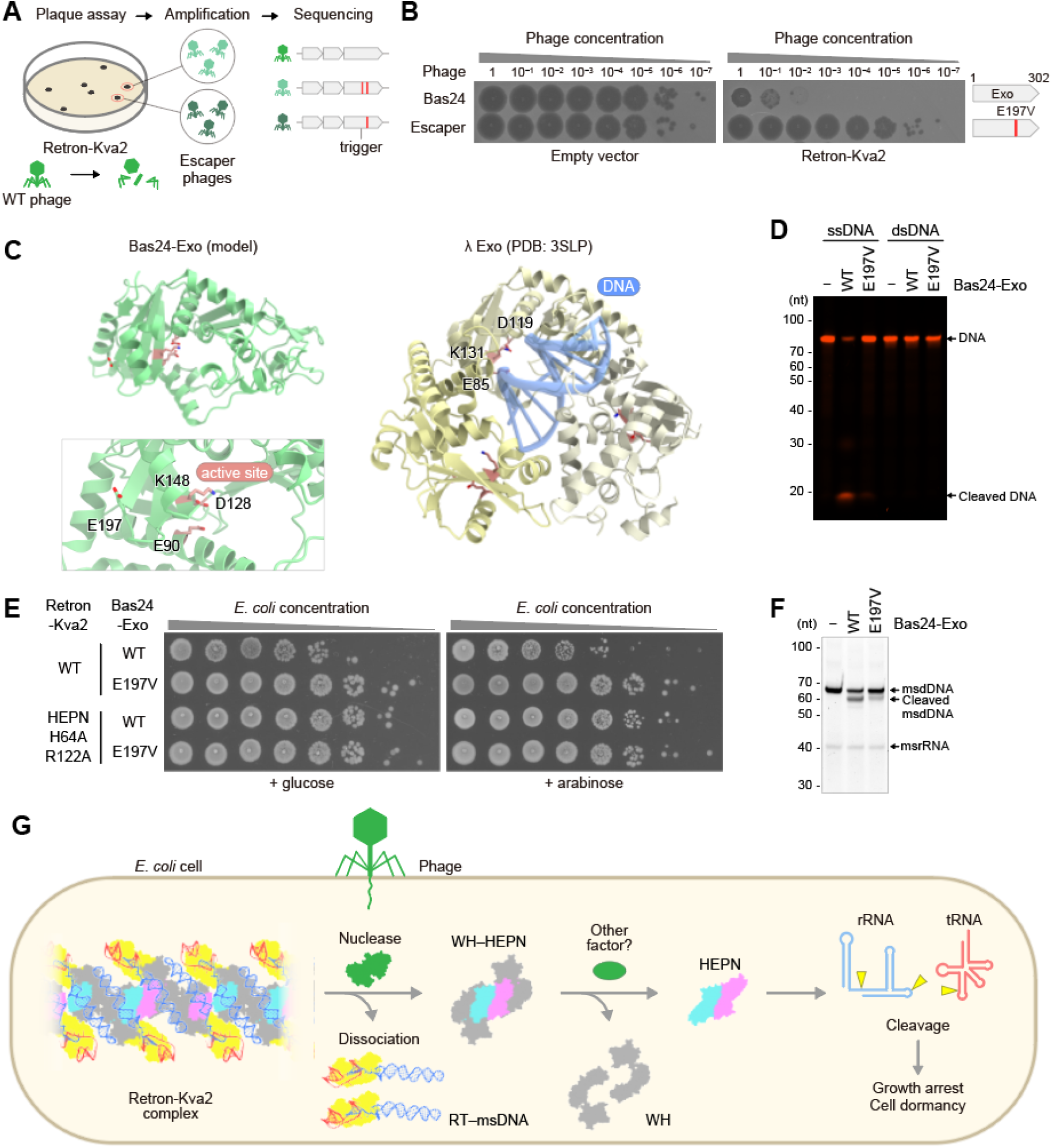
Activation of Retron-Kva2 by the phage nuclease. (**A**) Isolation process of the Retron-Kva2 escaper phages. (**B**) Spot assays of the isolated escaper Bas24 phages. *E. coli* cells harboring the Retron-Kva2 plasmid or an empty vector were infected with either the Bas24 phage or the escaper phage. The escaper phage carries the E197V mutation in a putative exonuclease (Bas24-Exo). (**C**) AlphaFold3-predicted model of Bas24-Exo. The crystal structure of λ exonuclease bound to double-stranded DNA (PDB: 3SLP) is shown on the right for comparison. While λ exonuclease forms a trimer, the oligomeric state of Bas24-Exo is unknown. The conserved catalytic residues and the E197V mutation are highlighted with stick models. (**D**) Nuclease activity of Bas24-Exo. Single- or double-stranded DNA substrates labeled with Cy5 at the 5′ end were incubated with purified WT Bas24-Exo or the Bas24-Exo E197V mutant at 37°C for 30 min, and then analyzed by denaturing urea-PAGE. The nucleic acids were visualized with fluorescent signals. Assays were repeated at least three times with similar results. (**E**) Growth of *E. coli* cells harboring plasmids encoding WT Retron-Kva2 or Retron-Kva2 with the HEPN mutations (H64A/R122A), along with plasmids encoding WT Bas24-Exo or Bas24-Exo with the E197V mutation, in the presence of 0.2% glucose or 0.2% arabinose. Growth defects were observed in the bacterial cells harboring plasmids encoding WT Retron-Kva2 and plasmids encoding WT Bas24-Exo, even in the presence of glucose, suggesting that the toxicity of Retron-Kva2 (HEPN effector) was not completely suppressed under this condition. Assays were repeated at least three times with similar results. (**F**) *In vitro* msdDNA cleavage by Bas24-Exo. The purified Retron-Kva2 complex was incubated with purified WT Bas24-Exo or its mutant (E197V) at 37°C for 30 min, and then analyzed by denaturing urea-PAGE. The nucleic acids were visualized with SYBR Gold. Assays were repeated at least three times with similar results. (**G**) Proposed model of the Retron-Kva2 anti-phage defense.

In other retron systems such as Retron-Eco7, phage-encoded nucleases cleave an msdDNA region within the RT–msDNA–effector complex to facilitate the release of the toxic effector, leading to abortive infection^28^. We therefore hypothesized that Bas24-Exo cleaves the msdDNA within the Retron-Kva2 complex, facilitating the dissociation of the HEPN–WH subcomplex. We incubated WT Bas24-Exo and the E197V mutant with the Retron-Kva2 complex (HEPN H64A) and examined msdDNA cleavage by urea-PAGE. WT Bas24-Exo cleaved the msdDNA, but not the msrRNA, within the Retron-Kva2 complex, whereas the E197V mutant failed to cleave the msdDNA efficiently (Figure 5F). Together, these results suggest that msdDNA cleavage by Bas24-Exo facilitates the dissociation of the HEPN–WH subcomplex from the Retron-Kva2 complex, thereby contributing to the activation of the Retron-Kva2 system (Figure 5G).

## Discussion

In this study, we showed that, in the type IX Retron-Kva2 system, the HEPN effector nuclease cleaves host RNAs, such as rRNAs and tRNAs, leading to growth arrest and conferring anti-phage defense. Furthermore, our cryo-EM analysis revealed that the RT–msDNA module and the WH ancillary effector interact with the HEPN nuclease to suppress HEPN-mediated RNA cleavage prior to phage infection. Thus, our findings support the concept that retron systems mediate anti-phage defense through toxin–antitoxin mechanisms, in which the cognate effectors serve as toxins and induce growth arrest in phage-infected bacterial cells, while the RT–msDNA module functions as an antitoxin, suppressing the effector’s toxicity prior to phage infection.

Unexpectedly, the cryo-EM structure of the Retron-Kva2 complex reveals that one RT–msDNA–HEPN–WH complex interacts with another complex to form a dimeric assembly, which further interacts with adjacent dimeric complexes through extensive contacts between the HEPN subunits (Figure 6). This unique interaction network leads to the formation of a sheet-like, supermolecular architecture distinct from previously reported retron complexes (Figure 6). While the Retron-Eco1 complex forms a supramolecular assembly^19,20^, substantial structural differences exist between Retron-Kva2 and Retron-Eco1. In Retron-Eco1, an RT–msDNA module and an *N*-glycosidase effector form a dimeric unit, primarily through effector–effector interactions, and the dimeric unit further associates with adjacent dimeric units through RT–RT interactions, thereby forming a filamentous assembly (Figure 6). The other three structurally characterized retron complexes—Retrons Eco2, Eco7, and Eco8—adopt distinct assemblies. The Retron-Eco2 RT–msDNA–TOPRIM effector nuclease complexes form a trimer^34,35^, while the Retron-Eco8 RT–msDNA–OLD effector nuclease assembles into a tetramer^30–33^ (Figure 6). In the Retron-Eco7 complex, two PtuA ATPase–PtuB HNH nuclease molecules associate with the RT–msDNA module, primarily through PtuA–msDNA interactions^23,25–29^ (Figure 6). As previously described^19,28^, the RT subunits from different retron systems share a right-hand-like core fold and commonly adopt the transcription-termination state, in which reverse transcription of the ncRNA (msdRNA region) is terminated at a specific position due to interactions between its thumb domain and the conserved stem loop in the ncRNA (msrRNA region). However, a structural comparison among the RT subunits from different retron systems also reveals distinct structural elements that contribute to the unique assemblies of the retron complexes, suggesting the co-evolution of RT and effector subunits in retron systems. For example, the N-terminal domain of the Retron-Kva2 RT subunit mediates essential interactions with the WH subunit. A structural comparison among these retron complexes also illuminates substantial differences in their msDNA architectures. In the Retron-Kva2, Retron-Eco1, and Retron-Eco7 complexes, the msDNAs contain a long, double-stranded DNA that plays a central role in complex formation (Figure 6). In contrast, in the Retron-Eco2 and Retron-Eco8 complexes, the msDNAs adopt a single-stranded conformation to mediate the symmetric assemblies of the complexes (trimer for Retron-Eco2 and tetramer for Retron-Eco8) (Figure 6).

**Figure 6.**
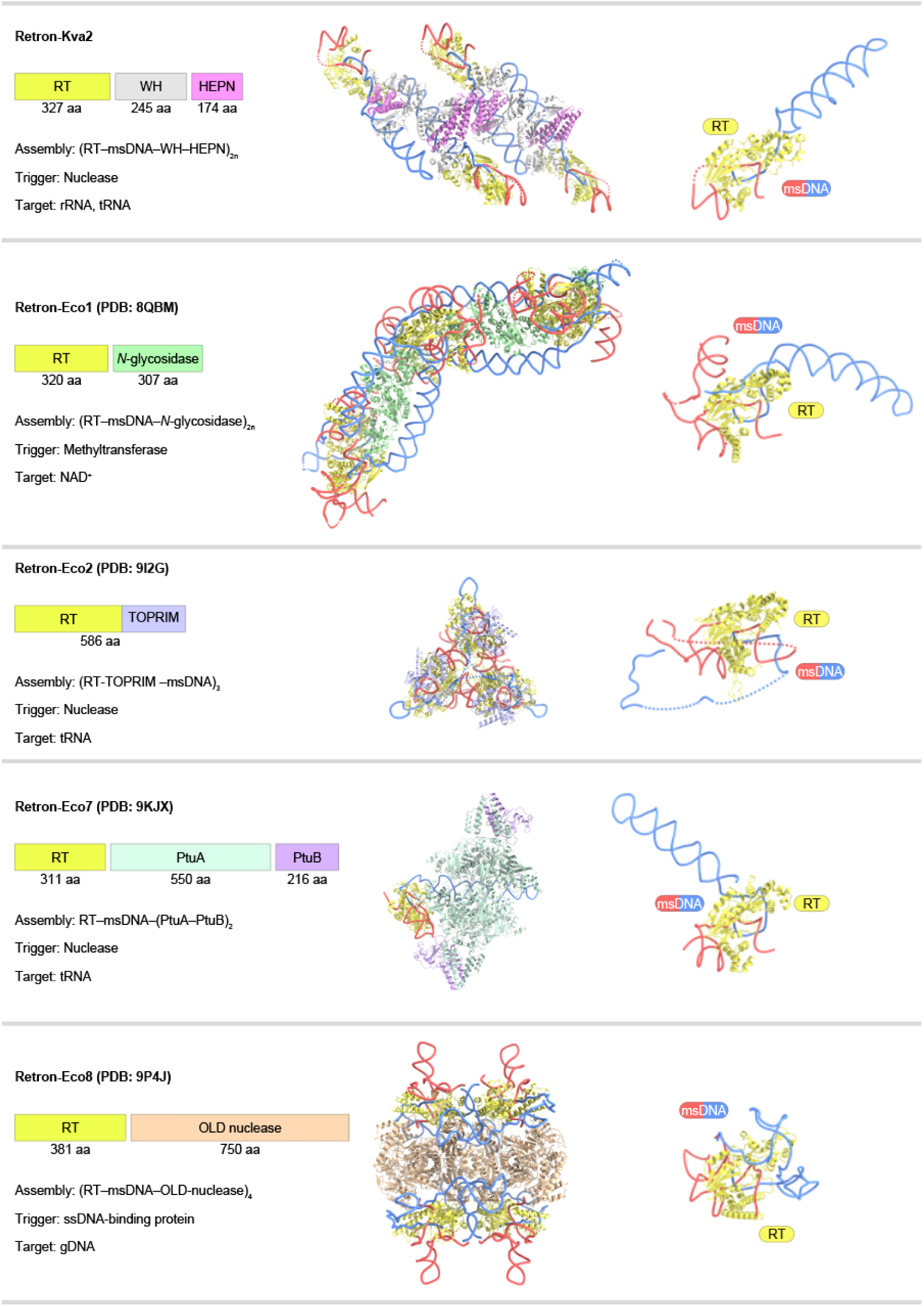
Comparisons of Retron-Kva2 with other retron complexes. The retron components, the RT–msDNA–effector complex structures, and the RT–msDNA subcomplex structures for Retron-Kva2, Retron-Eco1 (PDB: 8QBM), Retron-Eco2 (PDB: 9I2G), Retron-Eco7 (PDB: 9KJX), and Retron-Eco8 (PDB: 9P4J) are shown on the left, center, and right, respectively.

We identified the Bas24-Exo nuclease as one of the phage-encoded triggers that activate the Retron-Kva2 system, and demonstrated that Bas24-Exo cleaves the msdDNA within the Retron-Kva2 complex more efficiently than its E197V variant found in the escaper phages. These findings suggest that phage-encoded nucleases cleave the msdDNA scaffold in the Retron-Kva2 complex to facilitate the dissociation of the HEPN effector nuclease, leading to growth arrest in phage-infected cells, as observed in other retron systems, such as Eco2^34,35^ and Eco7^23,25–29^. However, the co-expression of Bas24-Exo was insufficient to fully activate Retron-Kva2, suggesting the requirement of other factors. Consistently, the expression of the HEPN effector alone caused severe growth defects, but its toxicity was attenuated by co-expression with the WH gene. In the Retron-Kva2 structure, the WH proteins extensively interact with the HEPN dimer, thereby restricting the access of RNA substrates to the HEPN active site. Together, our findings indicate that, unlike other known retron systems, the Retron-Kva2 system suppresses the toxicity of the HEPN effector using both the RT–msDNA module and the WH ancillary effector as dual antitoxins, and suggest that additional factors facilitate the dissociation of the HEPN toxin from the WH antitoxin.

We showed that the HEPN effector cleaves several host RNAs, including rRNAs and tRNAs, reinforcing the emerging theme that effector nucleases cleave the RNA components of the translational machinery to trigger abortive infection in various bacterial defense systems, including HEPN-MNT^37,40^, PARIS^46,47^, Retron-Eco7^24^, Retron-Eco2^35^, and CRISPR-Cas13^38,39^. Notably, diverse nucleases are responsible for the cleavage of distinct RNA targets in these defense systems. For example, the HNH nuclease specifically cleaves tRNA^Tyr^ in Retron-Eco7^24^, while the TOPRIM nuclease degrades various tRNAs in Retron-Eco2^35^. In contrast, the TOPRIM nuclease mediates the specific cleavage of tRNA^Lys^ in the PARIS system^46,47^. Intriguingly, previous studies reported that defense-associated HEPN nucleases display distinct substrate specificities: the CRISPR-Cas13 HEPN domain selectively cleaves a subset of tRNAs at their uridine-rich anticodons^38,39^, while the HEPN effectors in the HEPN-MNT systems from *Aphanizomenon flos-aquae*^37^ and *Legionella pneumophila*^40^ cleave subsets of tRNAs and rRNAs (both 23S and 16S rRNAs), respectively. Therefore, the Retron-Kva2 HEPN effector is unique in its capability to cleave both tRNAs and rRNAs, likely due to its distinct active-site architecture lacking the canonical Rx_4–6_H catalytic motif.

This study revealed that the type IX Retron-Kva2 system is activated upon phage infection, cleaving subsets of host rRNAs and tRNAs to induce growth arrest and facilitate anti-phage defense through abortive infection. Collectively, our findings highlight the functional and mechanistic diversity of retron-mediated immunity and enhance our understanding of various bacterial defense systems.

## Limitations of the Study

Our cryo-EM analysis revealed that the RT–msDNA module and the HEPN and WH effector proteins form a sheet-like supermolecular complex, and mutational data indicated that the formation of the dimeric units within the complex is essential for anti-phage defense. However, the functional role of the higher-order assembly remains unknown, particularly because the dimeric complex—the HEPN dimer bound to two RT–msDNA–WH subcomplexes—can form even in the absence of higher-order assembly. In addition, our data suggest that msdDNA cleavage promotes the release of the HEPN–WH subcomplex from the Retron-Kva2 complex, followed by the dissociation of the HEPN nuclease from the WH subunit; however, additional studies are required to confirm these molecular events. Although the Retron-Kva2 HEPN effector exhibits relaxed substrate specificity, cleaving both rRNAs and tRNAs, the mechanisms of its RNA recognition and cleavage remain unknown. Our biochemical data indicate that Bas24-Exo preferentially degrades single-stranded DNA and cleaves the msdDNA at a specific position, suggesting targeting of the msdDNA loop within the complex, but the precise cleavage site is unidentified. Further biochemical and structural work will be necessary to address these questions.

## Resource Availability

### Lead Contact

Requests for further information and resources should be directed to and will be fulfilled by the lead contact, Hiroshi Nishimasu.

### Materials Availability

All plasmids generated in this study are available from the lead contact with a completed materials transfer agreement.

### Data and code availability

The structure model of the Retron-Kva2 complex has been deposited in the Protein Data Bank, under the accession code 27LG. The cryo-EM density maps of the Retron-Kva2 complex have been deposited in the Electron Microscopy Data Bank, under the accession code EMD-81236. PDB IDs 3SLP, 8QBM, 9I2G, 9KJX, 9P4J, 9KIX, 7AE8, 5VRF, and 7XJG were used for structural comparisons.

## Acknowledgements

We thank the staff scientists at The University of Tokyo’s cryo-EM facility, especially Y. Sakamaki, for help with cryo-EM data collection. K.Yoneyama is supported by JSPS KAKENHI (Grant Number 26KJ0950). K.C. is supported by JSPS KAKENHI (Grant Number 25K18809), and Takeda Science Foundation (Medical Research Grant, Infectious Disease). M.Hiraizumi is supported by JSPS KAKENHI (Grant Numbers 26H01563 and 26K01944), JST ACT-X (Grant Number JPMJAX232F), and Takeda Medical Research Foundation. K.K. is supported by JSPS KAKENHI (Grant Numbers 25K21732 and 26H02321), AMED (Grant Numbers 26wm0325081, 26gm1610002, 26ae0121045, 26fk0108698, A519TR and S1-125502-00), and Shionogi Infectious Disease Research Promotion Foundation. H.N. is supported by the Platform Project for Supporting Drug Discovery and Life Science Research (Basis for Supporting Innovative Drug Discovery and Life Science Research (BINDS)) from AMED (Grant Number JP21am0101115; support number 6464), JSPS KAKENHI (Grant Numbers 21H05281 and 25H00436), JST CREST (Grant Number JPMJCR23B6), Takeda Medical Research Foundation, and the Inamori Research Institute for Science.

## Author Contributions

Y.H., Y.M., and K.Yoneyama performed functional and structural analyses with assistance from J.I., M.Hiraizumi, K.Yamashita, and H.N.; K.C. performed phage and sequencing experiments with assistance from M.Hashino, K.H., and K.K.; Y.H., Y.M., K.Yoneyama, and H.N. wrote the manuscript with input from all authors; K.K. and H.N. supervised the research.

## Declaration of generative AI and AI-assisted technologies in the writing process

During the preparation of this work, the authors used ChatGPT to check for grammar and spelling errors and improve the overall readability of the text. After using these tools, the authors reviewed and edited the content as needed and took full responsibility for the content of the publication.

## Methods

### Phage cultivation and spot assay

All phages used in this study were sourced from the BASEL collection and our in-house phage library, and are listed in Table S3. Spot assays were performed by mixing an overnight *E. coli* culture (DH10B or MG1655ΔRM) carrying the indicated plasmids with LB top agar (0.5% agarose and 1 mM CaCl_2_) and casting onto LB agar plates. Where indicated, 0.2% L-arabinose or 0.2% D-glucose was supplemented to the culture medium to induce or repress Retron-Kva2 expression, respectively. Serial 10-fold dilutions of phage stocks were prepared in SM buffer (50 mM Tris-HCl, pH 7.5, 100 mM NaCl, 5 mM MgSO_4_, and 0.01% (w/v) gelatin), and 2 µL of each dilution was spotted onto the bacterial lawn. Plates were incubated at room temperature until spots had been absorbed, followed by overnight incubation at 37°C, and plaques were scored the next day. Spot images were captured utilizing an EPSON GT-X980 flatbed scanner. The images presented in the figures are representative of three independent biological replicates. Where applicable, plaque-forming units (PFU) per mL were enumerated and plotted using GraphPad Prism 10.

### Liquid infection assay

To assess anti-phage defense in liquid culture, overnight *E. coli* ΔRM cultures harboring the indicated plasmids were diluted 1:100 into LB medium containing 1 mM CaCl_2_ and 20 µg/mL chloramphenicol. Aliquots (200 µL) were distributed into the wells of a 96-well plate, and phage solutions (4 µL) were added per well at multiplicity of infection (MOI) values of 0.1, 1, and 10. Wells including 4 µL of SM buffer were used as uninfected controls. Growth was monitored at 37°C with continuous shaking by measuring the OD_600_ at 10-min intervals over 6 h, using a SynergyNeo2 plate reader (Agilent Technologies). Raw data were analyzed and plotted using GraphPad Prism 10.

### Toxicity assay

Toxicity assays were performed as previously described^28^. Overnight bacterial cultures were serially diluted 10-fold in LB medium, and 4 µL of each dilution was spotted onto LB plates containing the appropriate antibiotics and supplemented with either 0.2% L-arabinose or 0.2% D-glucose. Plates were incubated at 37°C overnight, and colony formation was recorded using an EPSON GT-X980 flatbed scanner. All images shown are representative of three independent biological replicates.

### Total RNA extraction

For total RNA extraction from Retron-Kva2-expressing cells, *E. coli* DH10B cells containing the pLG-HEPN–WH–ncRNA–RT plasmid were grown in 1 mL of LB medium at 37°C for 3 h, and then infected with T5 phage (MOI = 1) or left uninfected (control) for 1.5 h. *E. coli* cells containing the empty pLG vector were used as the control. The cultures were then centrifuged at 6,000 g for 5 min, and the pellets were washed twice with buffer H (20 mM Tris-HCl, pH 8.0, and 150 mM NaCl). Total RNA was extracted using an miRNeasy Mini Kit (QIAGEN) following the manufacturer’s instructions. *E. coli* cells containing the pBAD-HEPN (WT, H64A, or R122A) plasmids were grown in 1 mL of LB medium at 37°C for 3 h. Expression was induced with 0.2% L-arabinose, and the cultures were incubated at 37°C for 1 h, with shaking at 200 rpm. *E. coli* cells containing the empty pBAD vector (pKLC23) were used as the control. The cultures were then centrifuged at 6,000 g for 5 min, and the pellets were washed twice with buffer H. Total RNA was extracted using an miRNeasy Mini Kit (QIAGEN). The extracted RNA samples were used for RNA-fragment sequencing.

### RNA-fragment sequencing

Total RNA samples were fractionated using urea-PAGE, and the RNA fragments were excised from the gel and extracted in 1× TBE buffer at 4°C overnight, followed by ethanol precipitation. For 3′ dephosphorylation and 5′ phosphorylation of the extracted RNA fragments, 800 ng of RNA fragments were incubated at 37°C for 1 h in a reaction mixture, containing 1 × T4 PNK reaction buffer, 20 U T4 polynucleotide kinase (New England Biolabs), and 20 U RNase Inhibitor, Murine (NEB). ATP was then added at a final concentration of 1 mM, and the mixture was further incubated at 37°C for 1 h, followed by phenol/chloroform/isoamyl alcohol (PCI) extraction and ethanol precipitation. NGS libraries were prepared using a NEBNext^®^ Multiplex Small RNA Library Prep Set for Illumina^®^ (New England Biolabs) according to the manufacturer’s instructions. Briefly, the workflow included 3′-adapter ligation, 5′-adapter ligation, cDNA synthesis, and PCR amplification. Size selection was performed using AMPure XP beads (Beckman Coulter) at a 1.8× bead-to-sample ratio to select for target fragments larger than 100 bp. Library quality was assessed using a MultiNA II MCE-301 microchip electrophoresis system (Shimadzu). Sequencing of the prepared library was performed by Plasmidsaurus Inc. RNA-fragment reads were processed on the Galaxy platform. Adapter sequences were trimmed from raw reads using Cutadapt^48^, and reads shorter than 20 nt after trimming were discarded. Processed reads were mapped to the *E. coli* DH10B reference genome (CP000948.1) using Bowtie^49^. Mapped reads were analyzed using bedtools^50^ for assignment to RNA features and extraction of FASTA sequences. Sequence logos showing preferences around cleavage sites were generated using WebLogo^51^.

### Construction of the *E. coli hupA::sfGFP* strain

The strain carrying an sfGFP insertion at the *hupA* locus was constructed by λ Red-mediated recombineering, following the established protocol^52^. Briefly, *E. coli* MG1655 ΔRM harboring pKD46 was grown overnight and subcultured (1:100) into 20 mL of fresh LB supplemented with 0.2% (w/v) arabinose. The culture was incubated at 28°C until it reached an OD_600_ of 0.5, then chilled on ice and harvested by centrifugation. Using ice-cold 10% (v/v) glycerol, the cell pellet was washed twice and finally resuspended in 200 μL. An 80-μL aliquot of this suspension was mixed with the purified PCR product and subjected to electroporation. The linear DNA fragment encoding the sfGFP-kanamycin-resistance cassette was amplified by PCR from pKCN-809, using primers bearing ∼40-nt 5′ extensions homologous to the 3′ end of *hupA*. Transformants were selected on LB agar containing kanamycin (50 μg/mL) and incubated overnight. Correct in-frame insertion was verified by colony PCR and sequencing.

### Live cell imaging

To visualize the nucleoid upon Retron-Kva2 HEPN overexpression, cells expressing a HupA::sfGFP fusion were imaged. Agarose pads were prepared by the slide-spacer method^53^. In brief, agarose was added to 25% (v/v) LB medium at a final concentration of 1% (w/v). The molten agarose was supplemented with carbenicillin (100 ng/µL), arabinose (0.1%, w/v), and FM4-64 (2 µg/mL; Invitrogen). The mixture was dispensed between two microscope slides separated by glass spacers and gently pressed to form a pad of uniform thickness, which was left to set at room temperature for 30 min and trimmed before use. Single colonies were collected from agar plates and suspended in 1 mL of 1× PBS, and 1–2 µL of each suspension was deposited onto a pad and incubated at 37°C for 2 h. As a positive control for nucleoid compaction, chloramphenicol (20 ng/µL) was applied onto the pad surface after the 2-h incubation. The pads were then inverted onto an ibidi µ-Slide 8 Well chamber for observation. Imaging was performed on a Zeiss LSM900 confocal microscope fitted with an Airyscan 2 detector and operated through the ZEN 3 software, using a 63× oil-immersion objective (NA 1.4) in the Airyscan super-resolution mode at a frame size of 2,048 × 2,048 pixels. sfGFP was excited at 488 nm (emission, 490–560 nm) and FM4-64 at 561 nm (emission, 561–700 nm). Each image was recorded in the mean-intensity mode with 8× line averaging. Airyscan reconstruction was performed with the default ZEN 3 parameters, which were kept constant across all samples within a given experiment.

### Sample preparation

For the purification of the Retron-Kva2 complex, the DNA fragment encoding HEPN (residues 1–174), WH (residues 1–245), ncRNA (173 nt), and RT (residues 1–327) was amplified by PCR and cloned into the pET28a vector to construct pET-His_6_-HEPN(H64A)–WH–ncRNA–RT-Twin-strep (HEPN with an N-terminal His_6_ tag and RT with a C-terminal Twin-strep tag). The gene encoding Bas24-Exo (residues 1–302) was cloned into a modified pE-SUMO vector with an N-terminal His_6_-SUMO tag (LifeSensors). To prepare the mutants, mutations were introduced using a PCR-based method, and the sequences were confirmed by DNA sequencing.

For the expression of the Retron-Kva2 complex, *E. coli* Rosetta 2(DE3) (Novagen) cells transformed with pET-His_6_-HEPN(H64A)–WH–ncRNA–RT-Twin-strep were cultured at 37°C until the OD_600_ reached 0.8, and protein expression was then induced with 0.2 mM IPTG (Nacalai Tesque). The *E. coli* cells were further cultured at 20°C overnight, harvested by centrifugation, resuspended in buffer A (20 mM Tris-HCl, pH 8.0, 300 mM NaCl, 3 mM 2-mercaptoethanol, and 1 mM PMSF), and lysed by sonication. The lysates were centrifuged, and the supernatant was applied to Strep-Tactin XT 4 Flow high-capacity resin (IBA). The complex was then eluted with buffer B (20 mM Tris-HCl, pH 8.0, 50 mM biotin, 300 mM NaCl, and 3 mM 2-mercaptoethanol). The complex was purified on a Superdex 200 Increase 10/300 GL column (Cytiva), equilibrated with buffer C (20 mM HEPES-NaOH, pH 7.5, 300 mM NaCl, and 1 mM DTT). The purified complex was stored at −80°C until use, except for the cryo-EM analysis, for which the freshly prepared complex was used without freezing. The protein concentration was measured using a Qubit Protein Assay Kit (Thermo Fisher Scientific).

For the expression of His_6_-SUMO-tagged Bas24-Exo, *E. coli* Rosetta 2(DE3) cells transformed with pE-SUMO-Bas24-Exo were cultured at 37°C in LB medium until the OD_600_ reached 0.8, and then induced with 0.2 mM IPTG and cultured overnight at 20°C. The cells were harvested, resuspended in buffer D (20 mM Tris-HCl, pH 8.0, 1 M NaCl, 20 mM imidazole, 3 mM 2-mercaptoethanol, and 1 mM PMSF), and lysed by sonication. The lysates were centrifuged, and the supernatant was applied to Ni-NTA Superflow resin (QIAGEN), which was washed with buffer E (20 mM Tris-HCl, pH 8.0, 300 mM NaCl, 20 mM imidazole, and 3 mM 2-mercaptoethanol), and eluted with buffer F (20 mM Tris-HCl, pH 8.0, 300 mM NaCl, 300 mM imidazole, and 3 mM 2-mercaptoethanol). After treatment with SUMO protease (prepared in-house), the protein was further purified on a HiTrap Heparin column (Cytiva), equilibrated with buffer G (20 mM HEPES-NaOH, pH 7.5, 150 mM NaCl, and 1 mM DTT). The protein was eluted with a linear gradient of 0.15–2 M NaCl. The purified protein was stored at −80°C until use.

### Sequencing of the msdDNA

For the purification of msdDNA, the Retron-Kva2 complex was purified using the Strep-Tactin XT 4 Flow resin, and then incubated at 37°C for 15 min with 1 U RNase A (New England Biolabs), 2 mM MgCl_2_, and 2 mM CaCl_2_. Proteinase K (Nacalai Tesque) was added, and the mixture was incubated at room temperature for 5 min, followed by PCI extraction and ethanol precipitation. NGS libraries were prepared from the purified msdDNA using an xGen^TM^ ssDNA & Low-Input DNA Library Prep Kit (IDT), according to the manufacturer’s instructions. Briefly, samples were denatured at 95°C for 2 min and immediately placed on ice. ssDNA-to-dsDNA conversion and adapter ligation were performed sequentially using the Adaptase^TM^ module, extension module, and ligation module. Size selection was performed using SPRISelect beads (Beckman Coulter) at a 1.2× bead-to-sample ratio for a 200 bp target fragment size. Libraries were indexed using xGen CDI Primers (IDT) and amplified by indexing PCR. Post-PCR cleanup was performed twice using SPRISelect beads at a 0.8× ratio to remove unincorporated primers, as required for sequencing on patterned flow cells. Library quality and size distribution were assessed using an Agilent TapeStation. Libraries were sequenced on an Illumina NextSeq 2000 platform (P2 cartridge, 300 cycles) in the paired-end mode (PE150). Illumina adapter sequences and Adaptase^TM^ tails were removed using cutadapt (v4.4). The following parameters were applied: 3′ adapter sequences AGATCGGAAGAGCACACGTCTGAACTCCAGTCAC (Read 1) and AGATCGGAAGAGCGTCGTGTAGGGAAAGAGTGT (Read 2) were trimmed. Poly-G sequences introduced by the 2-color chemistry of the NextSeq 2000 platform were trimmed using the --nextseq-trim option. Reads shorter than 15 bp after trimming were discarded (--minimum-length 15). Quality trimming was applied at a Phred score threshold of 20 (-q 20). Trimmed reads were aligned to a combined reference sequence comprising the *E. coli* BL21(DE3) genome (NCBI accession CP001509.3) and the retron expression plasmid using READemption2, which employs segemehl as the underlying aligner. Alignment was performed in the paired-end mode (--paired_end) with the following parameters: minimum read length 15 nt (--min_read_length 15), minimum Phred quality score 20 (--min_phred_score 20), maximum fragment length 500 bp (--max_fragment_length 500), and alignment accuracy 95% (segemehl default). Strand-specific coverage files (wiggle format) were generated using the reademption coverage subcommand, normalized to reads per million (RPM). Forward and reverse strand coverage was calculated separately.

### Cryo-EM analysis

The purified RT–msDNA–HEPN(H64A)–WH Retron-Kva2 complex solution (1.4 mg/mL) was mixed with 0.005% (final concentration) Tween 20 (Nacalai Tesque). The mixture (3 μL) was immediately applied to freshly glow-discharged Au 300 mesh R1.2/1.3 grids (Quantifoil), using a Vitrobot Mark IV (FEI) at 4°C, with a waiting time of 10 s and a blotting time of 6 s under 100% humidity conditions. The grids were plunge-frozen in liquid ethane cooled at liquid nitrogen temperature. Cryo-EM data were collected using a Titan Krios G3i microscope (Thermo Fisher Scientific), running at 300 kV and equipped with a Gatan Quantum-LS Energy Filter (GIF) and a Gatan K3 Summit direct electron detector in the correlated double sampling mode (The University of Tokyo, Japan). Movies were recorded at a nominal magnification of 105,000×, corresponding to a calibrated pixel size of 0.83 Å, with an electron exposure of 6.56 e^−^/pix/sec for 5.52 s, resulting in an accumulated exposure of 50.6 e^−^/Å^2^. The data were automatically acquired by the image shift method using the EPU software (Thermo Fisher Scientific), with a defocus range of −0.8 to −2.0 μm. The total dataset comprised 4,860 micrographs, including 4,588 acquired at 0° stage tilt and 272 acquired at 30° stage tilt.

### Image processing

Data processing was performed using the cryoSPARC v4.6.2 software package^54^. Dose-fractionated movies were aligned using Patch motion correction, and the contrast transfer function (CTF) parameters were approximated using Patch-Based CTF estimation with the default settings. Particles were automatically picked using template picker. The particles were curated by Heterogeneous Refinement, using the map derived from cryoSPARC Ab initio Reconstruction as the template. The curated particle set was refined using Non-uniform refinement with CTF value optimization and reference-based motion correction, yielding maps at 2.9-Å resolution, based on the Fourier shell correlation (FSC) = 0.143 criterion. To further improve the map quality in a monomeric unit, the particle set was refined using Local refinement with the focused mask, yielding local maps at 2.9-Å resolution. Local resolution was estimated using BlocRes in cryoSPARC^55^.

### Model building and validation

The model of the Retron-Kva2 complex was automatically built using Coot^56^, starting from a model built by ModelAngelo^57^. The model was refined using Servalcat^58^ against unsharpened half maps, with structural restraints from Boltz-2^59^ predictions processed by ProSMART^60^. The models were validated using MolProbity^61^. The statistics of the 3D reconstruction and model refinement are summarized in Table S1. The molecular graphics and cryo-EM density map figures were prepared using CueMol (http://www.cuemol.org) or UCSF ChimeraX^62^.

### Isolation of escaper phages

Overnight cultures of bacteria expressing Retron-Kva2 were diluted 1:100 into fresh LB medium containing 20 µg/mL chloramphenicol and 1 mM CaCl_2_, then incubated at 37°C for 30 min with shaking (200 rpm) before phages were introduced at an MOI of approximately 1. After overnight incubation, cultures were clarified by centrifugation at 5,000 g for 10 min and filtration through a 0.45 µm membrane. Serial dilutions (10-fold) of the filtrate were spotted onto a Retron-Kva2-expressing bacterial lawn and incubated at 37°C overnight. Four individual plaques were picked, propagated in 2 mL of the same medium with Retron-Kva2-expressing bacteria for 3 h, and clarified as described above. To confirm escape from Retron-Kva2 immunity, candidate escape mutants and wild-type phages were spotted onto lawns of bacteria carrying either an empty vector or a Retron-Kva2-encoding plasmid, prepared by embedding 300 µL of overnight culture in 10 mL of LB soft agar layered onto 1.5% LB agar with 20 µg/mL chloramphenicol, and incubated at 37°C overnight.

### DNA extraction and genome analysis of the escaper phages

High-titer lysates (>10^7^ PFU/mL) of mutant phages were prepared by PEG precipitation. Prior to precipitation, the filtered supernatant was treated with DNase I (Takara) and RNase A (Takara) at final concentrations of 3 U/mL and 20 ng/mL, respectively, and incubated at 37°C for 1 h. PEG8000 and NaCl were then added to final concentrations of 10% and 4% (w/v), respectively, and the mixture was incubated at 4°C overnight. After centrifugation (10,000 g, 1 h, 4°C), the resulting pellet was resuspended in 600 µL of SM buffer and mixed with 12 µL each of 10% (w/v) SDS and 0.5 M EDTA (pH 8.0), then heated at 68°C for 10 min. After cooling, 12 µL of 3 M sodium acetate (pH 6.5) was added, followed by the addition of 600 µL of phenol:chloroform:isoamyl alcohol (25:24:1). The mixture was vortexed for 30 s and centrifuged, and the aqueous phase was collected and extracted twice with 600 µL of chloroform:isoamyl alcohol (24:1) by vortexing and centrifugation at 16,000 g for 3 min. Phage DNA was precipitated by adding 2 volumes of ice-cold ethanol, 1/30 volume of 3 M sodium acetate (pH 6.5), and 1 µL of GlycoBlue^TM^ coprecipitant (Invitrogen, #AM9516), and incubating at −30°C for 30 min. The pellet was washed with ice-cold 75% ethanol, air-dried briefly at room temperature, and resuspended in 50 µL of sterile water.

Paired-end sequencing (2 × 150 bp) was performed on a NovaSeq 6000 platform (Azenta Life Sciences). Prior to assembly, adapter sequences were trimmed from raw reads using Cutadapt with a minimum read length of 15 bp. Wild-type phage genomes were assembled from trimmed reads using Shovill v1.1.0 with default parameters (https://github.com/tseemann/shovill), and gene annotation was performed on the assembled genomes using Pharokka v1.3.2. For variant calling, trimmed reads were subsampled to 100,000 read pairs per sample using seqtk (seed = 42) to ensure reproducibility and uniform coverage across samples. Single nucleotide polymorphisms (SNPs) were then identified by mapping the subsampled reads to the respective assembled reference genomes using Snippy v4.6.0 with default settings (https://github.com/tseemann/snippy). Large insertions or deletions were assessed manually by inspecting the resulting BAM files in Integrative Genomics Viewer (IGV) v2.16.0.

### *In vitro* DNA cleavage assay

To examine msDNA cleavage by Bas24-Exo, the purified Retron-Kva2 complex (200 nM) was incubated with the purified WT Bas24-Exo (500 nM) or its E197V mutant (500 nM) at 37°C for 30 min, in 10 μL of buffer I (20 mM Tris-HCl, pH 7.5, 150 mM NaCl, 7.5 mM MgCl_2_, and 1 mM DTT). To examine DNA cleavage by Bas24-Exo, the purified WT Bas24-Exo (100 nM) or its E197V mutant (100 nM) was incubated with single- or double-stranded DNA labeled with Cy5 at the 5′ end (200 nM) at 37°C for 30 min, in 10 μL of buffer I. The reactions were quenched with Proteinase K in denaturing buffer containing 7 M urea, and analyzed by 15% urea-PAGE. The gels were stained with SYBR Gold Nucleic Acid Gel Stain (Thermo Fisher Scientific), and then visualized using the FUSION Solo S imaging system (Vilber Bio Imaging).

