## Supplementary Information for "Structural mechanism of the type IX Retron-Kva2 anti-phage defense system"

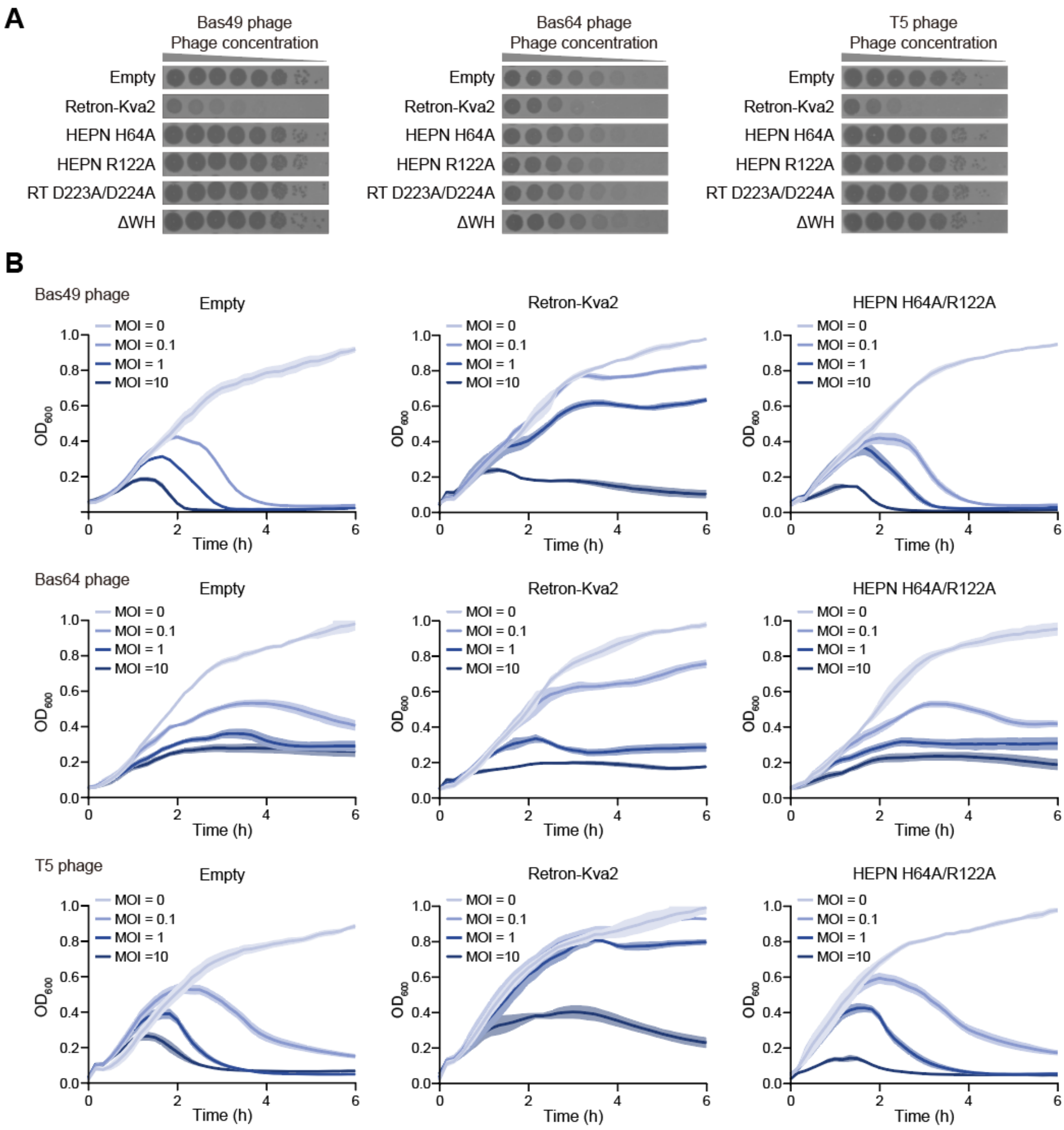


**Figure S1 | Anti-phage defense activity of Retron-Kva2.**

(**A**) Anti-phage defense activities of wild-type (WT) Retron-Kva2, Retron-Kva2 with the HEPN mutations (H64A or R122A), Retron-Kva2 with the RT mutations (D223A/D224A), and Retron-Kva2 with the WH deletion against the Bas49, Bas64, and T5 phages, evaluated by spot assays. *E. coli* cells transformed with an empty vector were used as the control. Assays were repeated at least three times with similar results.

(**B**) Growth curves of *E. coli* cells harboring an empty vector (−) or plasmids expressing Retron-Kva2 with or without the HEPN mutations (H64A/R122A). The cells were infected with either the Bas49, Bas64, or T5 phage at a multiplicity of infection (MOI) of 0, 0.1, 1, or 10. Data represent the mean ± SD of three biological replicates (n = 3).

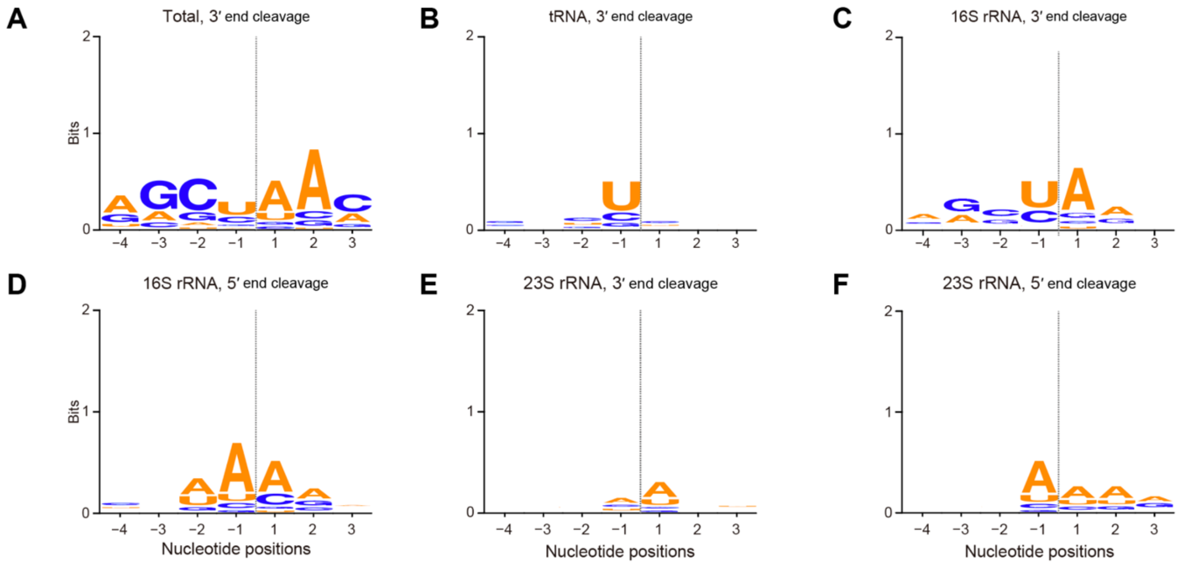


**Figure S2 | Nucleotide preference of Retron-Kva2 for RNA substrates.**

(**A**–**F**) Nucleotide enrichments around 3′ cleavage sites of total RNA cleavage products (**A**), tRNA 3′ cleavage sites (**B**), 16S rRNA 3′ cleavage sites (**C**), 16S rRNA 5′ cleavage sites (**D**), 23S rRNA 3′ cleavage sites (**E**), and 23S rRNA 5′ cleavage sites (**F**).

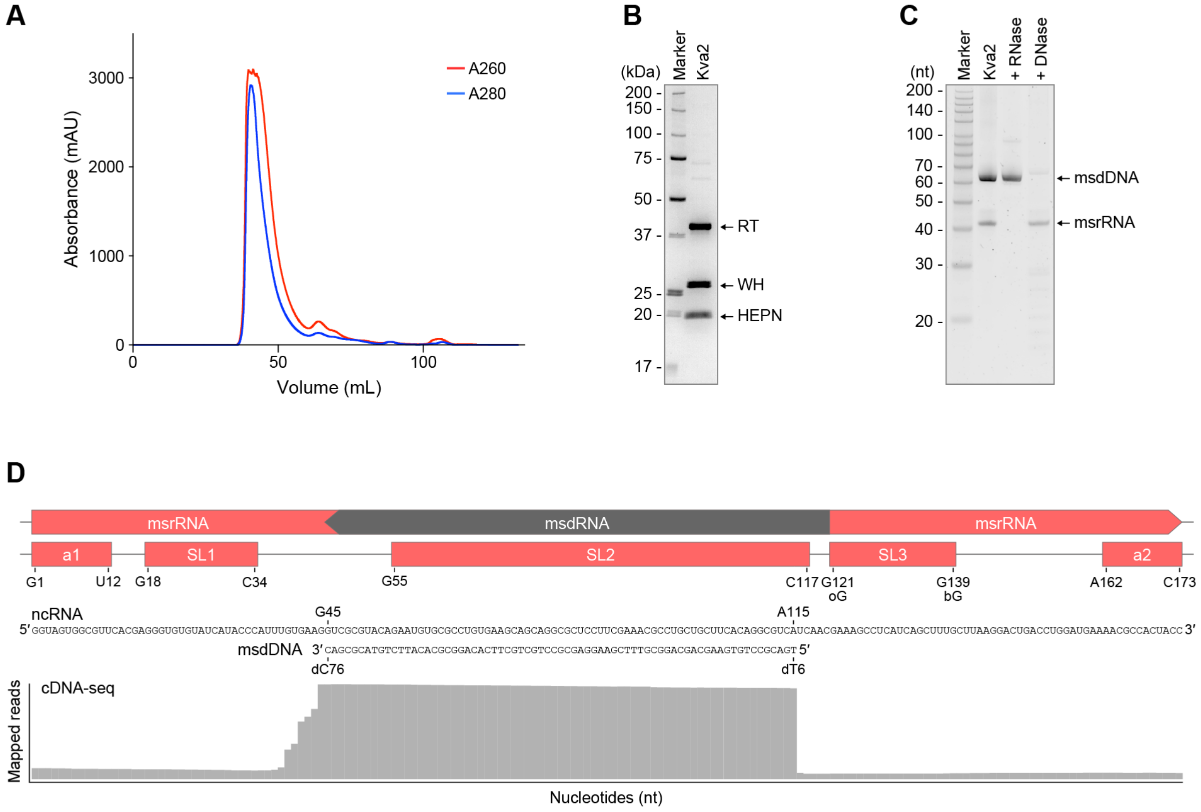


**Figure S3 | Preparation of the Retron-Kva2 complex.**

(**A**) Purification of the Retron-Kva2 (RT–msDNA–HEPN–WH) complex by size-exclusion chromatography.

(**B**) SDS-PAGE analysis of the purified Retron-Kva2 complex. Proteins were visualized with CBB staining.

(**C**) Urea-PAGE analysis of the purified Retron-Kva2 complex. The samples were treated with or without nucleases (RNase or DNase) and fractionated by urea-PAGE. Nucleic acids were visualized with SYBR Gold staining.

(**D**) Production of msdDNA from the ncRNA. DNA was extracted from the purified Retron-Kva2 complex and analyzed by NGS. Sequencing reads were mapped to the ncRNA region of the Retron-Kva2 expression plasmid. A schematic of predicted regions of the ncRNA is shown above the mapped reads.

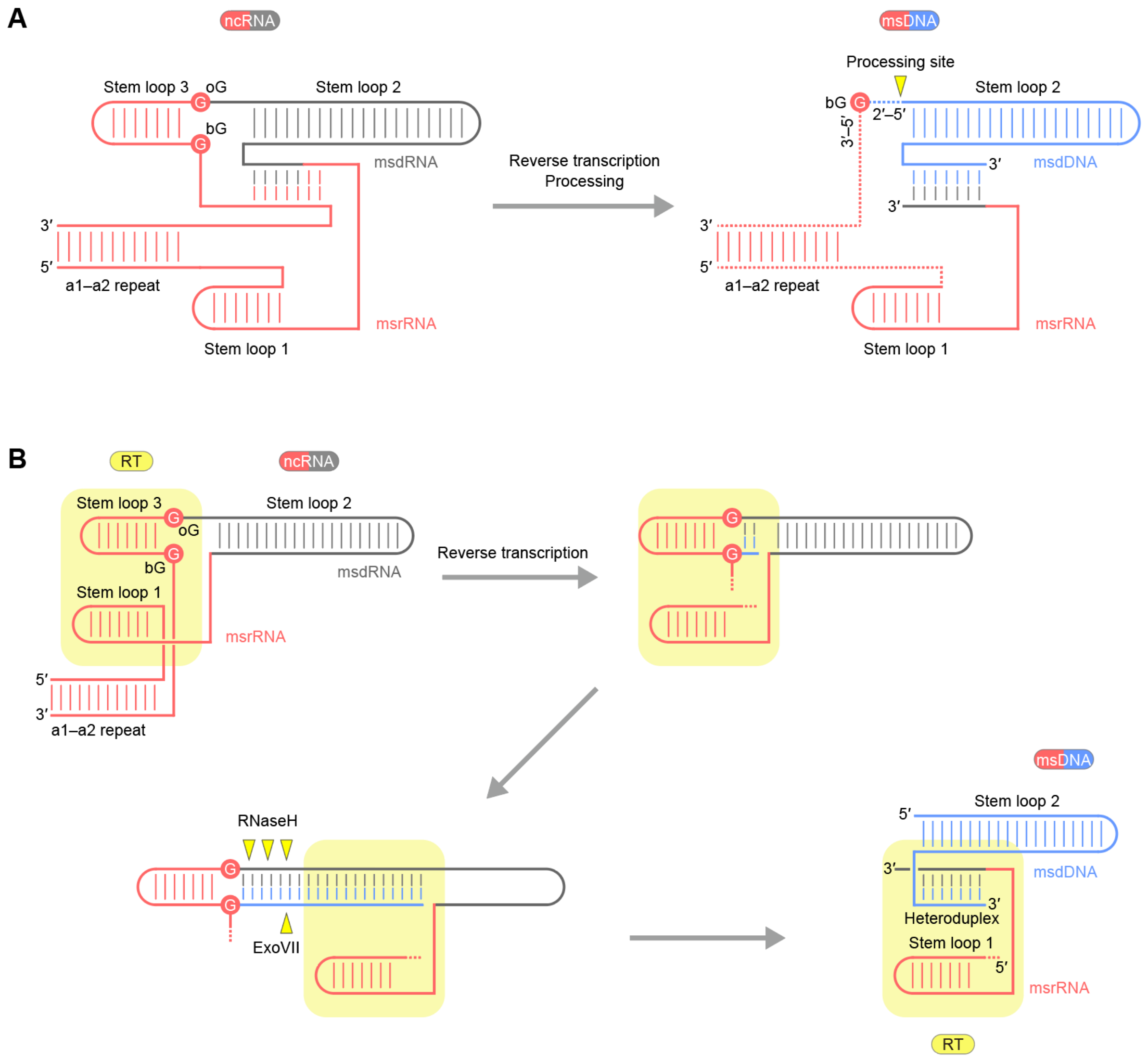


**Figure S4 | Retron-Kva2 msDNA synthesis.**

(**A**) Schematic of msDNA production in Retron-Kva2. bG, branching G; oG, opposing G.

(**B**) Schematic of the proposed reverse-transcription mechanism in Retron-Kva2.

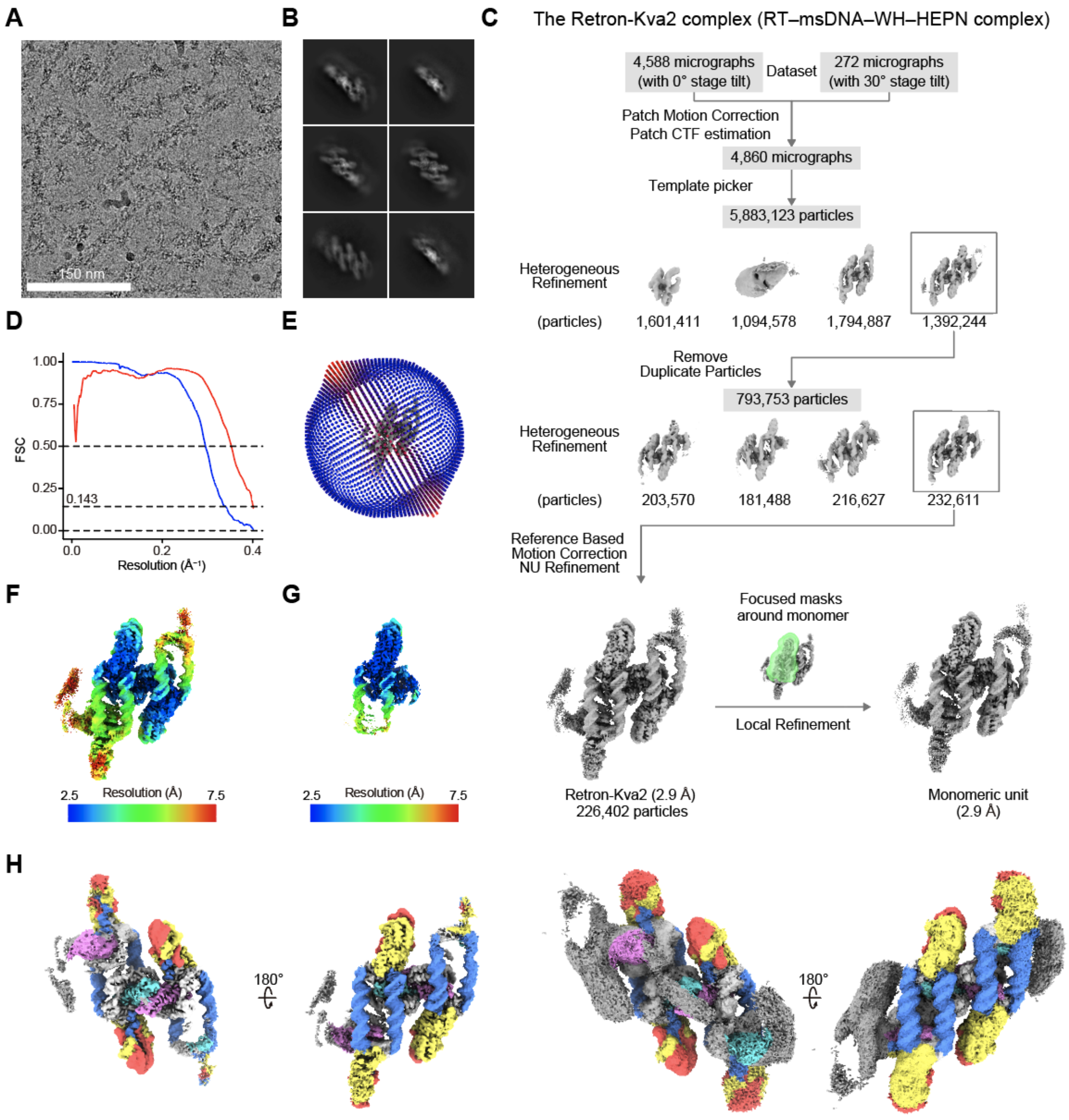


**Figure S5 | Cryo-EM analysis of the Retron-Kva2 complex.**

(**A**) Representative micrograph at a magnification of ×105,000.

(**B**) 2D averaged class images.

(**C**) Single-particle cryo-EM image processing workflow.

(**D**) FSC curves. The map-to-map FSC curves were calculated between the two independently refined half-maps after masking (blue line), and the overall resolution was determined by the gold standard FSC = 0.143 criterion. The map-to-model FSC curves were calculated between the refined atomic models and maps (red line).

(**E**) Angular distribution of particles in the final reconstruction.

(**F** and **G**) Cryo-EM density maps obtained by non-uniform refinement (**F**) and local refinement focused on the monomeric unit (**G**), colored according to the local resolution.

(**H**) Cryo-EM density maps at high (left) and low (right) contour levels, colored according to the protein domains.

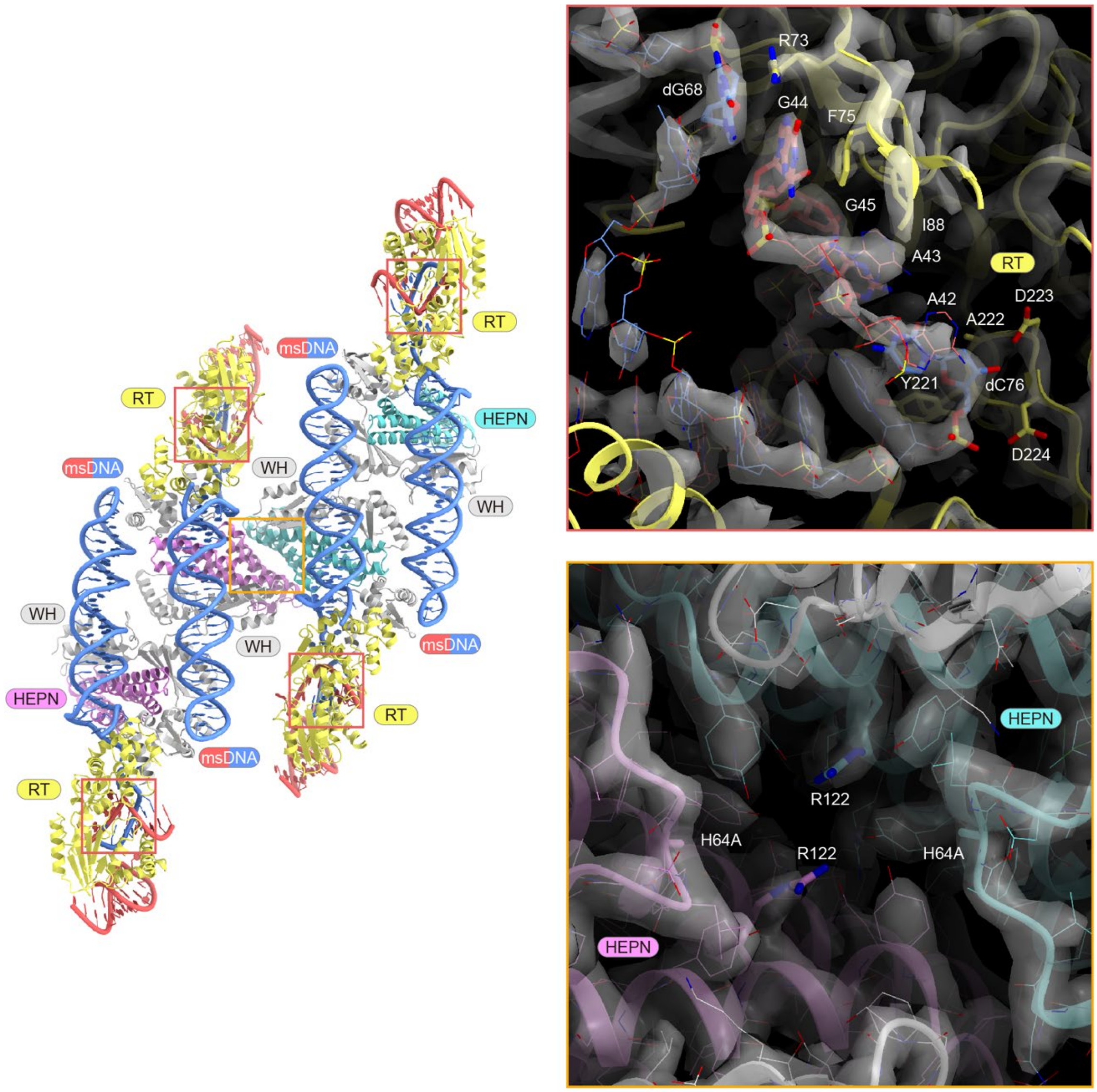


**Figure S6 | RT and HEPN active sites in the Retron-Kva2 complex.**

The RT and HEPN active sites are highlighted by red and orange boxes, respectively. Close-up views of the RT and HEPN active sites are shown on the right of the Retron-Kva2 complex. Key residues are shown as stick models, and cryo-EM density maps are displayed as semi-transparent gray surfaces.

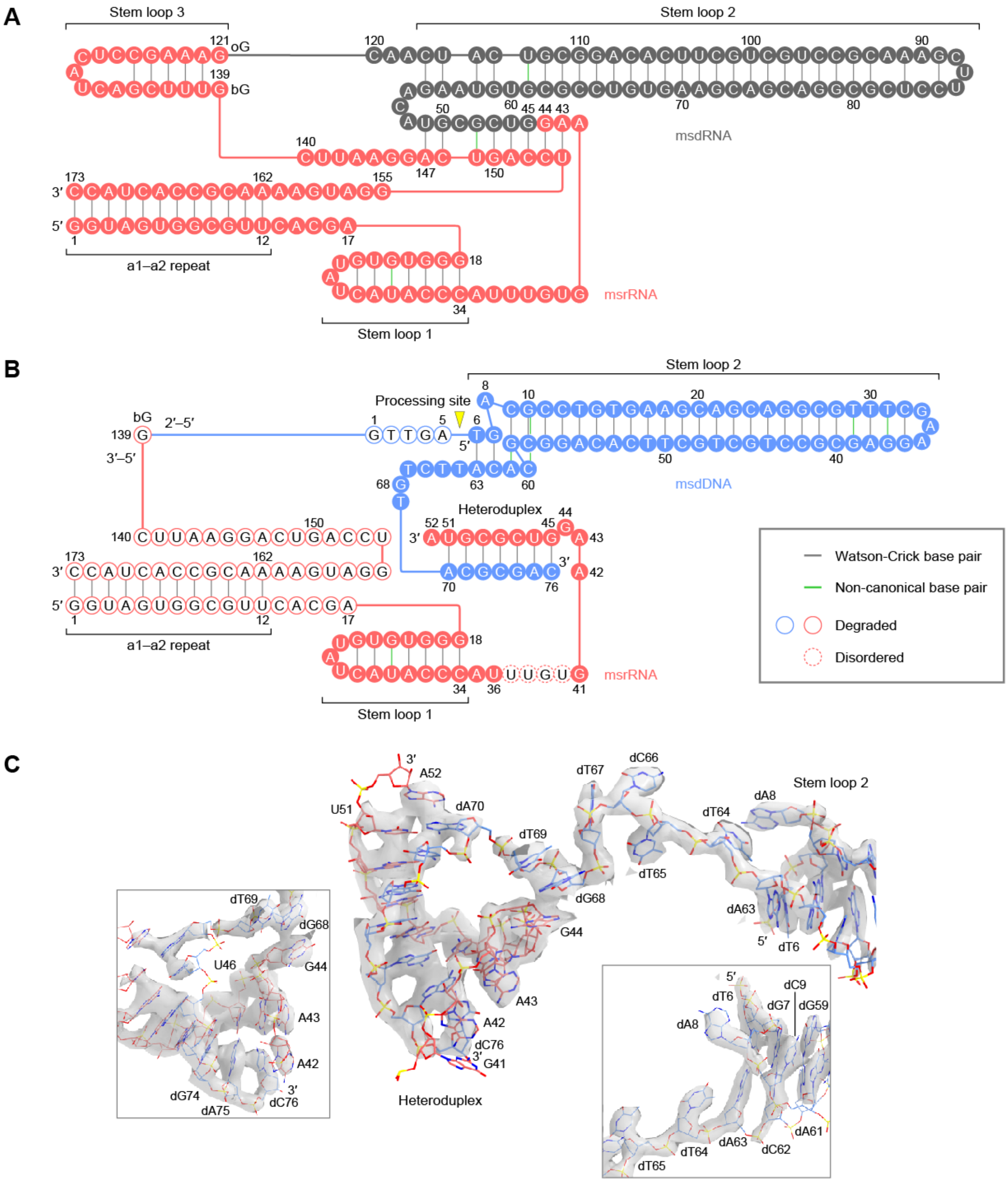


**Figure S7 | Nucleotide sequences of the ncRNA and msDNA.**

(**A** and **B**) Nucleotide sequences of the ncRNA (**A**) and msDNA (**B**) in the Retron-Kva2 system. Watson–Crick and non-canonical base pairs are indicated by gray and green lines, respectively. The site processed by ExoVII is marked by a yellow triangle. bG, branching G; oG, opposing G.

(**C**) Structures of the heteroduplex and stem loop 2 in the msDNA. Cryo-EM density maps are displayed as semi-transparent gray surfaces.

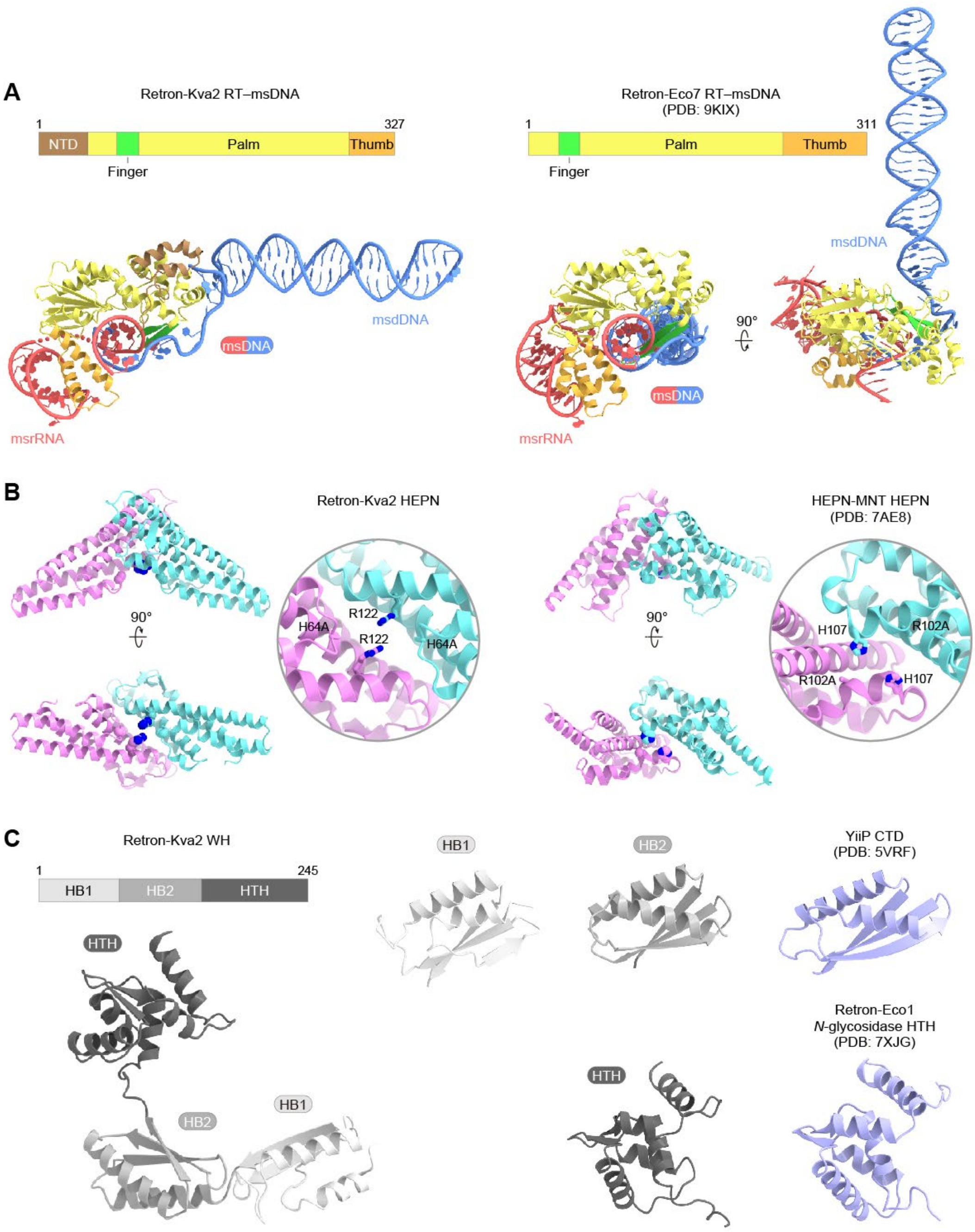


**Figure S8 | Structures of the RT, HEPN, and WH proteins in the Retron-Kva2 complex.**

(**A**) Structures of the RT–msDNA complexes from the Retron-Kva2 and Retron-Eco7 systems (PDB: 9KIX).

(**B**) Structural comparison of the Retron-Kva2 HEPN effector with the HEPN nuclease from the HEPN-MNT toxin–antitoxin system (PDB: 7AE8). Close-up views of the active sites are shown on the right of each structure.

(**C**) Structural comparison of the Retron-Kva2 WH protein with the C-terminal cytoplasmic domain of the zinc transporter (PDB: 5VRF) and the HTH domain of the *N*-glycosidase effector in the Retron-Eco1 system (PDB: 7XJG).

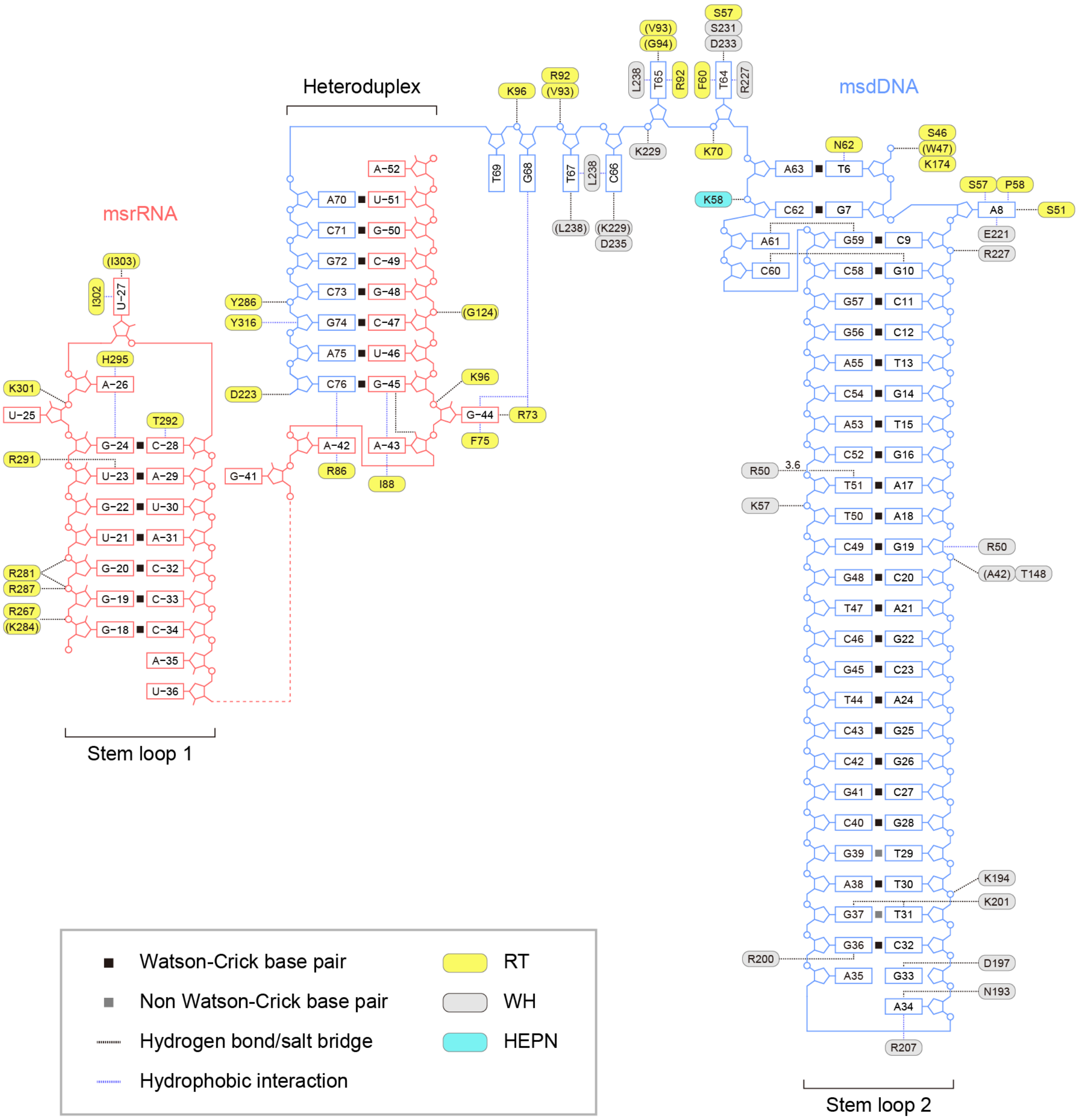


**Figure S9 | Schematic of msDNA recognition.**

The amino-acid residues that interact with the nucleic acids through their main chains are shown in parentheses.

**Table S1 | Cryo-EM data collection, refinement, and validation statistics.**

| | |
|---|---|
| **Data collection and processing** | |
| Microscope | Titan Krios G3i |
| Detector | K3 Camera |
| Automation software | EPU |
| Magnification | 105,000 |
| Voltage (kV) | 300 |
| Energy filter slit width (eV) | 25 |
| Exposure rate ($e^-/Å^2/s$) | 9.17 |
| Total electron exposure ($e^-/Å^2$) | 50.6 |
| Defocus range (μm) | −0.8 to −2.0 |
| Pixel size (Å) | 0.83 |
| Number of frames per image | 50 |
| Number of micrographs | 4,860 |
| Symmetry imposed | *C1* |
| Tilt angle (°) | 0 and 30 |
| Initial particle images (no.) | 5,883,123 |
| Final particle images (no.) | 226,402 |
| Map resolution (Å) | 2.9 |
| FSC threshold | 0.143 |
| **Model building and refinement** | |
| Model composition | |
| Protein residues | 2,960 |
| RNA residues | 124 |
| DNA residues | 284 |
| Protein atoms | 23,824 |
| RNA atoms | 2,656 |
| DNA atoms | 5,832 |
| Average B factors ($Å^2$) | |
| Protein | 372.7 |
| DNA | 460.4 |
| RNA | 383.4 |
| R.M.S. deviations from ideal | |
| Bond lengths (Å) | 0.0072 |
| Bond angles (°) | 1.1 |
| **Validation** | |
| Clashscore | 0.62 |
| Rotamer outlier (%) | 2.65 |
| Ramachandran plot | |
| Favored (%) | 96.76 |
| Allowed (%) | 2.83 |
| Outlier (%) | 0.41 |
